# An active electronic, high-density epidural paddle array for chronic spinal cord neuromodulation

**DOI:** 10.1101/2024.05.29.596250

**Authors:** Samuel R. Parker, Jonathan S. Calvert, Radu Darie, Jaeson Jang, Lakshmi Narasimhan Govindarajan, Keith Angelino, Girish Chitnis, Yohannes Iyassu, Elias Shaaya, Jared S. Fridley, Thomas Serre, David A. Borton, Bryan L. McLaughlin

## Abstract

**Objective:** Epidural electrical stimulation (EES) has shown promise as both a clinical therapy and research tool for studying nervous system function. However, available clinical EES paddles are limited to using a small number of contacts due to the burden of wires necessary to connect each contact to the therapeutic delivery device, limiting the treatment area or density of epidural electrode arrays. We aimed to eliminate this burden using advanced on-paddle electronics.

**Approach:** We developed a smart EES paddle with a 60-electrode programmable array, addressable using an active electronic multiplexer embedded within the electrode paddle body. The electronics are sealed in novel, ultra-low profile hermetic packaging. We conducted extensive reliability testing on the novel array, including a battery of ISO 10993-1 biocompatibility tests and determination of the hermetic package leak rate. We then evaluated the EES device in vivo, placed on the epidural surface of the ovine lumbosacral spinal cord for 15 months.

**Main results:** The active paddle array performed nominally when implanted in sheep for over 15 months and no device-related malfunctions were observed. The onboard multiplexer enabled bespoke electrode arrangements across, and within, experimental sessions. We identified stereotyped responses to stimulation in lower extremity musculature, and examined local field potential responses to EES using high-density recording bipoles. Finally, spatial electrode encoding enabled machine learning models to accurately perform EES parameter inference for unseen stimulation electrodes, reducing the need for extensive training data in future deep models.

**Significance:** We report the development and chronic large animal *in vivo* evaluation of a high-density EES paddle array containing active electronics. Our results provide a foundation for more advanced computation and processing to be integrated directly into devices implanted at the neural interface, opening new avenues for the study of nervous system function and new therapies to treat neural injury and dysfunction.

## Introduction

Epidural electrical stimulation (EES) of the spinal cord has been used for the treatment of neuropathic pain for almost six decades (Nahm 2020), with over 50,000 procedures performed in the United States each year (Krog et al. 2023). In therapeutic or research applications, EES leads are surgically inserted into the epidural potential space on the dorsal aspect of the spinal cord and connected to an internal or external pulse generator (IPG or EPG, respectively) using inline contacts. Recently, EES has been used in preclinical research settings to study central, peripheral, autonomic, and sensorimotor function (Gad et al. 2014; Parker et al. 2013; Calvert et al. 2023; Capogrosso et al. 2016; Musienko et al. 2012), as well as in clinical research settings to restore sensorimotor function following a spinal cord injury or amputation (Lorach et al. 2023; Rowald et al. 2022; Chandrasekaran et al. 2020; Nanivadekar et al. 2023). In parallel, epidural spinal recordings obtained through EES leads have been used as control signals for pain therapies (Nijhuis et al. 2023), to study voluntary movement control (Burke et al. 2021), and to examine somatosensory evoked spinal potentials (SEPs) (Nainzadeh et al. 1988; Woodington et al. 2024). However, all prior research studies and clinical applications utilize passive leads, requiring a one-to-one connection between electrical contacts on the tissue and the stimulation electronics. In practice, this has put a constraint on the number of contacts used to a maximum of 32 channels, resulting in limited selectivity during stimulation and resolution during recording.

To ensure proper placement of the EES electrodes for activation of targeted spinal circuits, a key step is intraoperative testing during implantation (Falowski 2019). During intraoperative testing, electrical stimuli are delivered through the implanted paddle. The stimuli primarily activate sensory neurons, which then recruit motor neurons via reflex pathways (Capogrosso et al. 2013). Using bilateral electromyography (EMG), the response of muscles at the same spinal level as the sensory targets is evaluated, and lead position is adjusted to achieve the desired placement (Shils and Arle 2018). To recruit more specific pools of neurons on the spinal cord, multipolar stimulation is applied in a current steering approach (Mishra, Kulkarni, and Gadgil 2023; Rowald et al. 2022; Lorach et al. 2023; Chandrasekaran et al. 2020; Nanivadekar et al. 2023), which requires access to multiple contacts in a confined region of interest. Subsequently, there have been efforts to design an EES paddle optimized for particular applications (for example, locomotor restoration), using neuroimaging data and computational modeling to guide the arrangements of contacts on the paddle (Rowald et al. 2022). Additionally, standard EES paddles principally deployed for the management of neuropathic pain concentrate their electrodes on the central structures of the spinal cord, and do not place stimulating contacts over lateral regions (for example, spinal nerve entry points or dorsal root ganglia (DRGs)). Activation of these structures has resulted in strong motor recruitment of target muscles (Soloukey et al. 2021), which may be more selective than stimulation more proximal to contralateral motor pools. However, the distribution of afferent entry points and DRGs is not uniform along the rostrocaudal axis, again highlighting the importance of proper intraoperative alignment when using sparsely distributed electrode contacts.

The necessity for manual stimulation parameter selection is a major barrier to the widespread adoption of EES for functional improvement after spinal cord injury (Solinsky, Specker-Sullivan, and Wexler 2020). Consequently, there have been efforts to automate the optimization of stimulation parameters using machine learning models (Govindarajan et al. 2022; Zhao et al. 2021). Although these approaches have shown improvement in speed, prior models were conditioned on each electrode independently. Such an approach is effective for paddles containing low numbers of contacts but requires significant computing power to run many models in parallel for high-count paddles. Additionally, conditioning on each electrode independently does not allow the model to infer what may happen during stimulation on electrodes excluded from the training set. As the number of stimulation contacts in a paddle increases, the time to collect training data scales linearly. Instead, if neural networks can infer the consequences of stimulation at unseen stimulation contacts, collecting training data from all contacts may not be necessary to rapidly program EES on many-contact paddles.

To address both placement limitations and stimulation parameter search optimization, a next-generation EES paddle must (1) span multiple spinal segments to target multiple sensorimotor pools, (2) provide a wide mediolateral span to target lateral structures, (3) enable localized current steering and bipolar re-referencing with densely packed electrodes,and (4) be reconfigurable for patient anatomy. These requirements are incompatible with conventional EES hardware with limited electrode contacts.

Thus, we designed a smart-implant called HD64 to provide 60 electrodes of epidural current steering for EES across a 14.5 mm mediolateral span (almost 2x wider than commercial EES paddles) and 2.5 vertebral segments (40 mm). Our smart-implant is only 2 mm thick, and sports a high-density electrode array integrated with a hermetic electronic multiplexing package and a 24:64 reconfigurable multiplexer ASIC - thus breaking the one-to-one wiring constraint of percutaneous EES hardware. The programmable smart-paddle enables the spatial layout of stimulating electrodes to be software-controlled, and is powered by a +/−5V AC power driver ASIC with fail-safe AC power leakage-detection circuits. For future human use of the device, we developed a good manufacturing practice (GMP) manufacturing process and performed ISO 14708 aging testing as well as ISO 10993-1:2021 biocompatibility testing. A comparison between three commercial EES paddles with the work presented here can be found in Table 1.

**Table 1.** A comparison of the physical characteristics of commercially available epidural electrical stimulation paddles and the paddle developed in this work, the HD64.

|  | **5-6-5** (Medtronic 2022) | **CoverEdge / CoverEdgeX** (Boston Scientific 2020) | **HD64 (this work)** |
| --- | --- | --- | --- |
| **Manufacturer** | Medtronic (Minneapolis, MN) | Boston Scientific (Marlborough, MA) | Micro-Leads Medical (Somerville, MA) |
| **Length** | 64.2mm | 50mm / 67mm | 57.5mm |
| **Treatment Width** | 10mm | 8mm / 8mm | 14.5 mm |
| **Thickness** | 2.0 mm | 2.0 mm | 0.2 mm edge, 2.0 middle |
| **Number of contacts** | 16 | 32 | 60 |
| **Inter-electrode spacing (mediolateral, rostrocaudal)** | (1.0mm, 4.5mm) | (1.0mm, 1.0mm) / (1.0mm, 3.2mm) | (0.7mm, 0.9mm) |
| **Density vs 5-6-5** | 1 | 2.1 / 1.4 | 2.2 |
| **Percutaneous lead-tails** | 2 | 4 | 2 |
| **Contact surface area** | 6mm<sup>2</sup> | 3.4mm<sup>2</sup> | 4.15mm<sup>2</sup> |
| **Contains an active multiplexer** | No | No | Yes |

The culmination of this work resulted in the long-term evaluation of the smart HD64 paddle for EES using benchtop verification and in-vivo validation in two sheep with up to 2 paddles per animal for 15 months. During this time, our goals were to establish the utility of HD64 to perform high density EES studies and observe any device-related functional issues. We quantified the selectivity of EES-evoked motor responses delivered using dense bipoles, differences in SEPs recorded using high- and low-density bipoles, and we extended the state-of-the-art stimulation parameter inference machine learning models to high-density electrodes (Govindarajan et al. 2022).

## Methods

### Treatment Span of Sensory-Motor EES

In the T9 to T12 thoracic region, the transverse diameter of the human spinal cord is 8-9 mm, and the epidural width is approximately 13-15 mm (Ko et al. 2004). Recent work has highlighted the benefit of activation of lateral structures, including DRGs, and their potential as a target for locomotor neuroprosthesis following spinal cord injury (Soloukey et al. 2021). Conventional surgical paddles for pain are limited to dorsal column medio-lateral treatment (<7.5 mm with 4 electrode columns) across two vertebral segments (<49 mm with 8 electrodes rows). To extend the medio-lateral therapeutic span of EES delivery, there is a need to access the dorsal horn and dorsal roots. Based on the desired treatment area, the paddle requirements are to treat a 13.7 x 43 mm stimulation area. Extending the treatment surface area for EES faces two challenges: (1) the lateral epidural volume adjacent to nerve roots is much thinner than at the midline, and (2) performing EES with a sparser electrode array would reduce the targeting resolution, potentially introducing off-target effects or non-specific activation.

### Mechanical requirements for surgical introduction of the lead

The paddle geometry was designed to mimic conventional EES paddles at the anatomical midline using a 2 mm thick geometry. Due to the 1 mm lateral epidural thickness, the paddle lateral edges were limited to 0.7 mm. Ten cross-sectional geometries of electrode profiles were developed in groups of three and sequentially evaluated. Each design was modeled using computer-aided design (CAD) software and molded in silicone to exhibit various longitudinal and lateral flexibility profiles. The profile groups were sent to three neurosurgeons to evaluate the preferred paddle design (criteria for their evaluation included steerability, axial ‘spine’ stiffness, and bendability) while accessing the epidural space through a small laminectomy. The resulting profile (Fig. 1l) was the final profile for two key reasons: (1) the lateral wings remained flexible and could bend through a conventional-width laminectomy, and (2) a rigid paddle spine was strongly preferred as it enabled physicians to advance the paddle rostrally using forceps.

**Figure 1.**
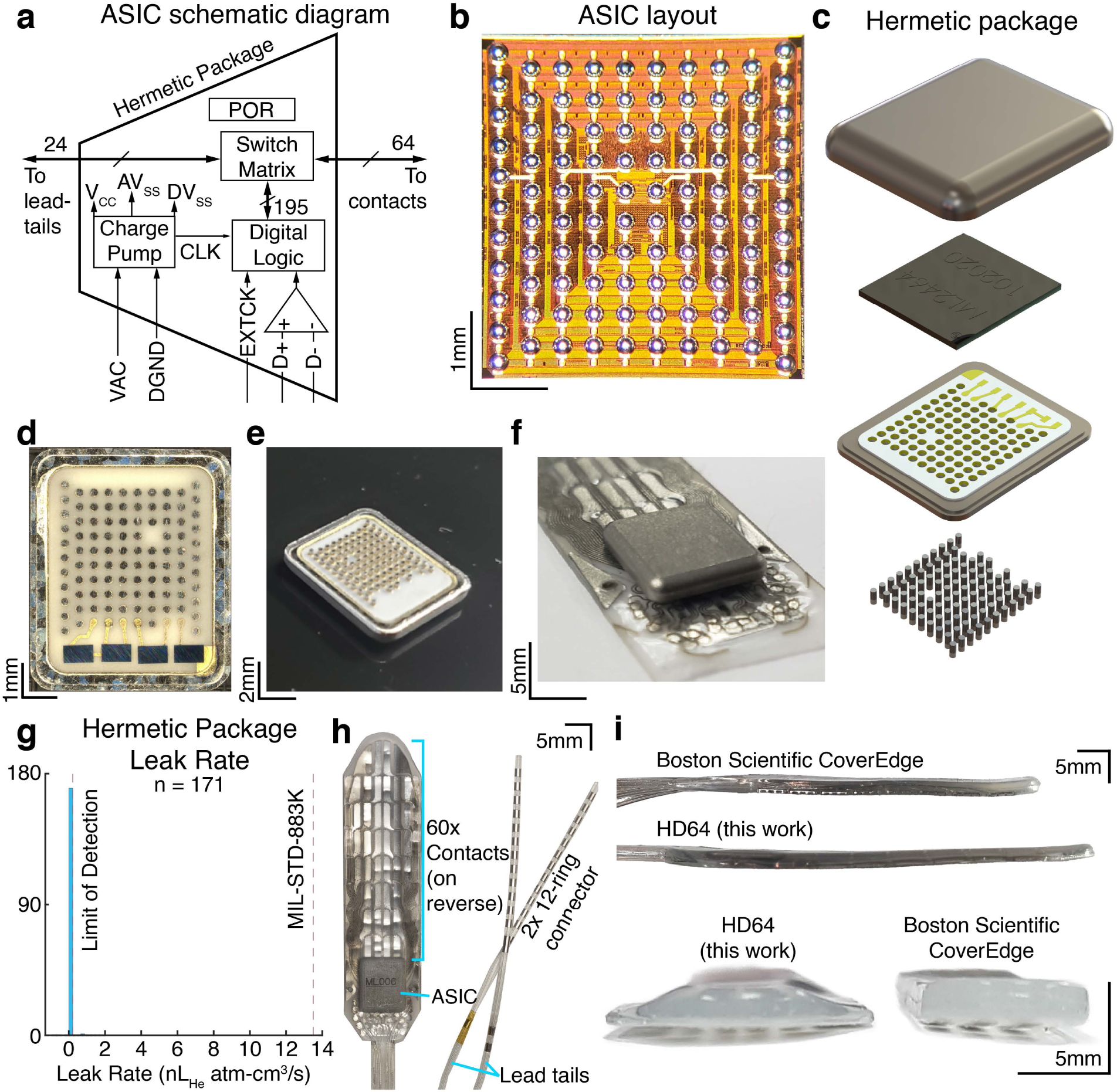
The HD64: Active electronics hermetically sealed into an epidural electrode paddle array. **a)** A functional block diagram of the application specific circuit (ASIC) embedded on the spinal paddle. **b)** a photograph of the ASIC die (redistribution layer and solder ball placement shown). **c)** An exploded 3D render of the hermetic package. From top to bottom, we show the Titanium 6-4 top case, the balled ASIC, a novel Alumina feedthrough array, and the feedthrough pins. **d, e)** Photographs of the assembled hermetic feedthrough package. **f)** The hermetic package is precision aligned within <10 μm onto the electrode before conductors are thermally micro-bonded followed by silicone injection molding. **g)** A histogram of the hermeticity leak rates measured from 171 hermetic packages. The MIL-STD-883K helium leak failure threshold is overlaid as the red dashed line. **h)** A photograph of the molded HD64 paddle array. **i)** Side- and front-on photographs comparing the lower-volume profile of the HD64 compared to a Boston Scientific CoverEdge32 surgical paddle.

### Smart-implant multiplexing architecture

Using a fixed number of electrodes (e.g., 16 or 32) spread across a wider medio-lateral span would result in a sparser electrode density and result in a reduced EES targeting resolution. Given the lateral nerve root density, a sparse density was undesirable. Conventional implanted surgical paddles have reached 32 electrodes, but do not exhibit stimulation sites over the nerve roots and require management and connection of 4 lead tails below the skin. Scaling to 64 electrodes using current connector technology would require 8 lead-tails and 64-channel connector on the IPG, which would greatly increase bulk in the body and surgical complexity. Since EES generally needs access to many electrodes but only a subset are used at the same time, we developed a multiplexing architecture to programmably connect electrodes to the pulse generator. Specifically, we sought to develop a scalable manufacturing process to produce paddle electrodes.

### Implant-grade design requirements

HD64 is a smart-paddle containing embedded electronics that is powered and controlled by a pulse generator over a two-lead tail wiring scheme. To realize an active-multiplexed electrode array, we developed multiple new implant technologies including: 1) a high-voltage multiplexer that operates from charge-balanced AC power (Fig. 1a, b), 2) a high-density and low-cost 93-pin hermetic feedthrough package with >80% yield (Fig. 1c-e), 3) precision electrode technology (platinum-iridium with 100 μm lines and 100 μm spaces), and 4) medical-grade micro-bonding processes to permanently bond 85 electrical connections between the hermetic electronic package and electrode array.

For long-term implanted use, we developed requirements based on our clinical inputs for people with spinal cord injury and/or chronic pain in the lower extremities. We established electrical and mechanical safety requirements and testing paradigms in accordance with ISO 14708-3:2017. We further developed a biocompatibility evaluation plan in accordance with ISO 10993-1:2018. The device also had to be manufactured according to GMP and undergo Ethylene Oxide (ETO) sterilization validation.

#### Electrical Safety: AC-powered implanted satellite devices

In this work, we describe two specific and critical design elements that enable new smart implants including: (1) ultra-low current satellite multiplexer operation using charge-balanced AC power over an implanted lead and pluggable-connector interface, (2) digital charged-balance programmable control of the distal satellite multiplexer. We tested commercial leads and connectors with +/−10V AC power using an accelerated aging paradigm in a sealed saline container. At the end of a 5 year implanted life equivalent, the saline was tested for residual traces of platinum-iridium using inductively coupled mass spectrometry (ICMS) to test for any electromigration. We determined that +/−5V AC through the lead-wire with a 2x voltage multiplier within the hermetic electronics package prevented any risk of electromigration. Additionally, we developed a separate IPG-side driver ASIC chip to deliver low-current +/−5V AC power with charged-balanced differential data lines to program the multiplexer. Board logic-level programming of a satellite multiplexer over a lead wire is not an acceptable risk, as any DC potential on the line can lead to electromigration at the connector. The driver ASIC was designed to deliver a programmable AC-current limited to 10 μA - 100 μA and 10 kHz to 500 kHz. Importantly, a novel real-time leakage current detection circuit was developed such that any broken wire or connection could be detected by the ASIC which would then generate an internal digital fault flag.

#### Biocompatibility evaluation

For translation to future human implanted use, we determined that the following tests must be performed: cytotoxicity, sensitization, irritation, material-mediated pyrogenicity, acute systemic toxicity, subacute toxicity, implantation, genotoxicity, ethylene oxide residuals, and partial chemical characterization. To achieve these endpoints, approximately 300 active-paddle test articles were sent to a certified ISO 10993-1 test house over a 14-month duration. All tests successfully passed these tests and no histological abnormalities were identified by the test house during any of these tests. A summary of the biocompatibility tests performed and their results are included in Supplementary Table 1.

#### Precision electrode technology

Precision electrodes electrodes were developed with a highly-novel delamination-free process using medical-implant grade silicone (Nusil, Carpentaria, CA), platinum-iridium 90/10 conductors, and a nylon mesh reinforcement layer (Fig. 1f). In this process, the silicone and platinum-iridium are pre-processed using a proprietary laser patterning technique, resulting in 50-100 μm electrode features in 50 μm thick platinum-iridium 90/10. A flexible reinforcement mesh layer was embedded within the silicone to prevent stretching of the silicone to protect the electrode traces from fracture during stretching or repetitive pull cycles. The HD64 paddle consists of three silicone and two metal layers for a total substrate thickness of 400 μm. Using a proprietary process, the silicone layers are chemically fused together to form a seamless and delamination-free construction for long-term implanted operation.

#### High-voltage 24:64 multiplexing ASIC with AC power and charge-balanced programming

An onboard multiplexer application-specific integrated circuit (ASIC) (Fig. 1a) was developed to support two key functions needed for this architecture: (1) a fail-safe low-voltage and low-current AC powering scheme with on-chip +/−9V compliance voltage multipliers and a bidirectional charge-balanced digital read-write scheme, (2) a 24:64 switch matrix to allow 24 bidirectional wires to connect to any of the 60 high-density electrodes. Fabrication was performed using the X-FAB XH035 18V 350 nm silicon process to construct the ASIC. A polyimide redistribution layer was developed to provide a solder-ball flip-chip interface between the ASIC (Fig. 1b) and hermetic feedthrough (Fig. 1c). The multiplexer is designed to be dynamically programmed using a +/−1.8V charge-balanced, differential digital interface operating from +/−5V AC power. Once powered, programming can be used to connect any of 24 input wires to 64 outputs. The multiplexer contains 64 blocks, each containing four switches to ensure every output electrode has a redundant connection to at least four stimulation/recording inputs. Switches were assigned to blocks in such a way as to maximize the number of switches that may become nonfunctional while still enabling sequential programming to raster through the entire 60-electrode array. The ASIC operates from AC power (+/−5V AC, 10-500 kHz) and features real-time current-limiting. Charge balance is necessary to prevent corrosion and long-term electro-migration of the conductors. The ASIC uses an on-board rectifier with off-chip capacitors to convert the AC signal to +/−9V DC internally and 3.3V.

#### High-density hermetic electronics package

We developed a custom 93-feedthrough ceramic assembly (8 x 8 x 0.75 mm) brazed into a titanium flange. On the interior of the package, the ASIC was flip-chipped to the surface of the feedthrough array (Fig. 1c). A low-outgassing underfill was applied between the ASIC and the ceramic and inspected for voids. A moisture getter was applied to the interior titanium lid, which was then seam-welded to form a hermetic assembly. Surface mount capacitors were soldered to metal pads on the ceramic feedthrough surface. The lid was attached to the feedthrough array and a laser seam-weld was performed in a nitrogen-helium environment. On the exterior of the hermetic packaging, the feedthroughs emerge from the ceramic surface and serve as a surface for 93 subsequent micro-bonds to be performed with the electrode array (Fig. 1d,e).

#### Electrode-hermetic electronics integration

The completed hermetic electronics assembly and high-density electrode assembly were then micro-bonded using a proprietary process. Rigorous machine vision was used to perform automated inspection of the 93 feedthroughs and receiving electrode pad positions. Components with >35 μm of tolerance were discarded to ensure the reliability of the process. Fixturing and vision cameras were used to align the components together until the feedthroughs and receiving pads were overlapping with <20 μm offset (Fig. 1f). Machine vision was used to inspect the tolerances of the electrode, and the feedthrough component and components with >35 μm of tolerance were discarded to ensure reliability of the bonding process. After the precision alignment was performed, a custom-made machine was developed to perform thermal welding between each platinum-iridium pad and each platinum-iridium feedthrough.

#### Injection molding and lead-tail attachment

The proximal end of the HD64 smart paddle uses two connector tails, each with 12 in-line ring contacts (Fig. 1h), which have decades of long-term implant reliability (Letechipia et al. 1991) and are compatible with IPGs. The lead-tails were manufactured and welded to the platinum-iridium receiving pads on the HD64 substrate. A silicone injection molding step was used to create the contoured paddle profile of the HD64, as well as to completely insulate between all of the feedthrough pins. The high-pressure injection molding process with appropriate ports and runners ensured that no air voids were present between feedthrough pins. All fixtures and processes were developed to ensure the silicone flow did not trap bubbles during the assembly process. Following this process, the HD64 was completely assembled and ready for use (Fig. 1h).

### Benchtop evaluation of the HD64

#### Hermeticity of the electronics package

To protect the embedded active electronics from the ionic environment in the epidural space, it is necessary to ensure the hermeticity of the electronics package and feedthrough assembly. Hermeticity model calculations were performed using a 2-year shelf life and a 1-year implanted life. Hermetic tests for electronics (MIL-STD-883 Method 1014.15) have failure limits based on the dew point inside a free volume package. The internal free volume of the HD64 assembly was 0.02 cm^3^, which corresponds to a failure leakage rate of 5×10^−9^ atm-cm^3^/sec of air. This is equivalent to 13.5×10^−9^ atm-cm^3^/sec of leakage using helium, which is used to test hermetic components. Using the laser interferometer method, described in the standard, the leak rate was measured for 171 hermetic packages. Fig. 1g shows the distribution of measured leak rate values. Less than 10% of assembled packages had gross leak rates and were sent for subsequent testing. All accepted packages recorded a leak rate at least 15 times lower than the failure threshold set by MIL-STD-883. Based on the empirical leak rates 5×10^−10^, hermeticity calculations suggest the internal free volume of the package will not reach a dew point until far beyond a 2-year shelf life and 2 year implanted use. We were therefore confident that the hermetic package could continue safely to *in vivo* testing.

#### Control of active-multiplexing

Establishing digital control and maintaining knowledge of the state of the HD64 multiplexer was necessary to demultiplex the recorded spinal responses and stimulation information *post-hoc*. A schematic representation of the connections and devices used in this manuscript is presented in Fig. 2a. The HD64 is powered and controlled by an external controller (MB-Controller, Micro-Leads Medical, Somerville, MA), which communicates with a host PC over a serial connection. MATLAB (version 2023b, MathWorks, Natick, MA) functions were provided to connect to, configure, and read from the multiplexer. Stimulation was also controlled using MATLAB. A custom script was written to ensure that the multiplexed stimulation channel was connected to the target electrode contact. Additionally, changes to the multiplexer configuration were logged alongside stimulation information, allowing synchronization with the recorded electrophysiological signals. This enabled recorded multiplexed signals to be split as the multiplexer configuration changed and a sparse 60 x *n* matrix of demultiplexed signals to be created, where *n* is the recording length in samples.

**Figure 2.**
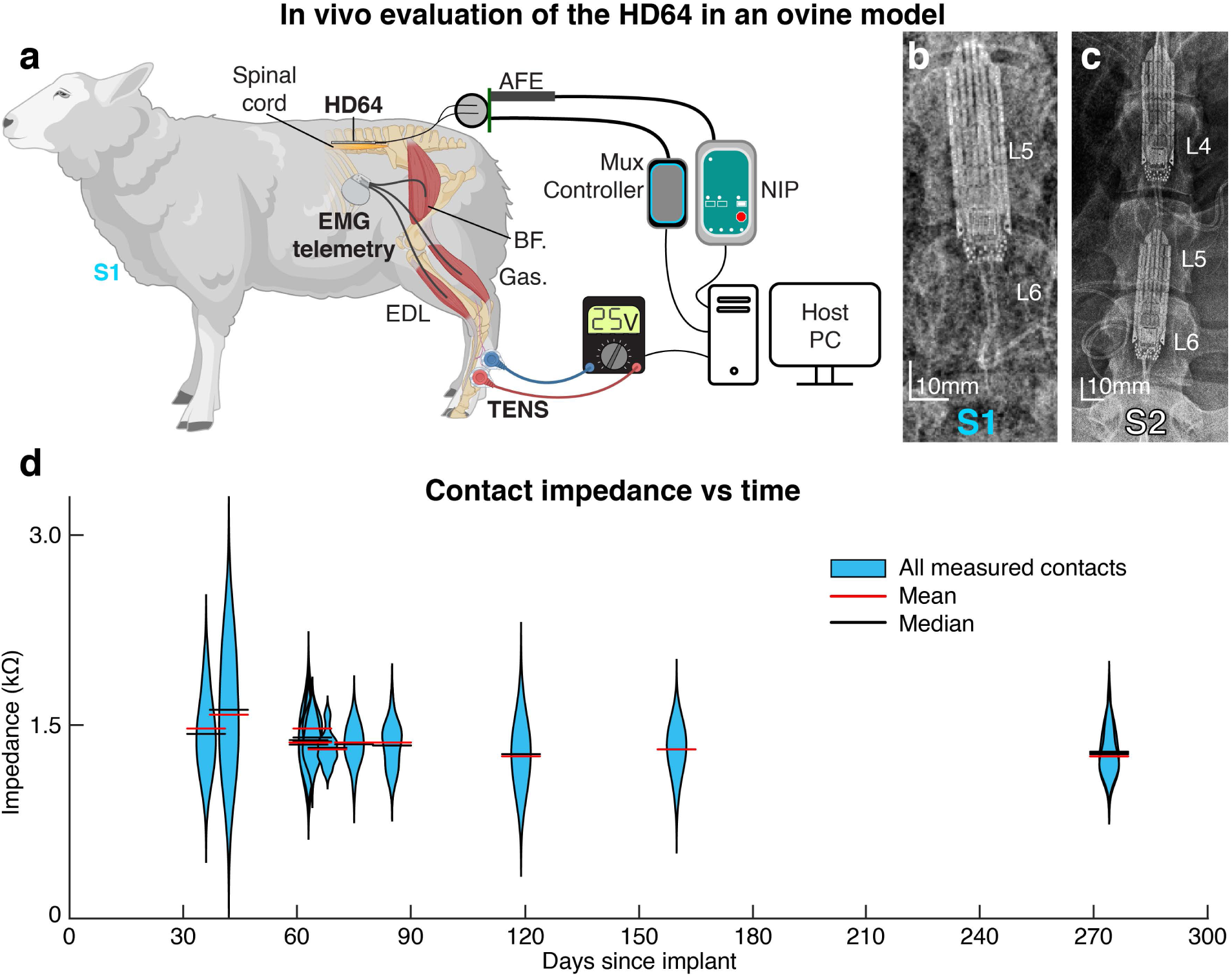
An overview of the *in vivo* evaluation of the HD64. **a)** A schematic representation of the connections and devices used to evaluate the HD64. The host PC controls the collection of spinal electrophysiology, application of transcutaneous electrical nerve stimulation (TENS) and epidural electrical stimulation (EES), and the configuration of the onboard ASIC through the multiplexer controller. In S1 and S2, electromyographic (EMG) signals are collected from implanted, wireless intramuscular EMG telemetry devices or wireless surface EMG devices, respectively. AFE is the analog front end, BF is the biceps femoris, Gas is the gastrocnemius, and EDL is the extensor digitorum longus. **b)** A radiograph of Sheep 1, showing a single HD64 implanted. **c)** A radiograph of Sheep 2, showing two HD64s implanted. **d)** The violin plots demonstrate that contact impedance remained stable over time, up to 274 days post-implantation.

### *In vivo* evaluation of the HD64

All surgical and animal handling procedures were completed with approval from the Brown University Institutional Animal Care and Use Committee (IACUC), the Providence VA Medical System IACUC, and in accordance with the National Institutes of Health Guidelines for Animal Research (Guide for the Care and Use of Laboratory Animals). Two sheep (both female, aged 4.19±0.3 years, weighing 92.5±2.5 kg) were used for this study (Fig. 2b,c). Recordings were performed with both sheep for 15 months. Animals were kept in separate cages in a controlled environment on a 12-hour light/dark cycle with *ad libitum* access to water and were fed twice daily. The ovine model was chosen for this study as the spine and spinal cord are comparable in size and share many anatomical features with humans, and the use of the ovine model to study the spinal cord has been well established (Wilson et al. 2017; Parker et al. 2013; Parker, Laird-Wah, and Cousins 2018; Parker et al. 2020; Reddy et al. 2018; Marcus et al. 1997). A visual overview of the experimental setup is shown in Fig. 2a.

#### Surgical procedures

The sheep were implanted, as previously reported (Calvert et al. 2023). However, S1 was implanted with a single paddle due to a narrow epidural space at L3. Briefly, under propofol-based general total intravenous anesthesia (TIVA), an L4 – L6 laminectomy with medial facetectomy was performed. The rostral HD64 paddle (if included) was gently placed, then slid rostrally, after which the caudal paddle was placed. Strain relief loops were made, and then the lead-tails were tunneled to the skin, where they were externalized. Reference and ground electrodes (Cooner AS636 wire, Cooner Wire Company, Chatsworth, CA) were secured epidurally and in the paraspinal muscles, respectively. Strain relief loops were made, and these wires were also tunneled then externalized. Intraoperative testing was used to confirm device functionality. The animals were allowed to recover from anesthesia then returned to their pens.

A second surgical procedure was performed to place intramuscular EMG recording equipment (L03, Data Sciences International, St. Paul, MN) in the lower extremity musculature of S1. The sheep was intubated and placed prone on the operating table. The sheep was placed in a V-shaped foam block to maintain stability throughout the procedure, and a rectangular foam block was placed under the hip to alleviate pressure on the hindlegs (Calvert et al. 2023). The legs were hung laterally off of the operating table, and each of the hooves was placed in a sterile surgical glove and wrapped in a sterile bandage for manipulation during the procedure. On each side, the *extensor digitorum longus*, *biceps femoris*, and *gastrocnemius* were identified by palpating anatomic landmarks. Small incisions were made over the bellies of each of the three muscles to be instrumented, and a small subcutaneous pocket was made on the flank. All three channels were tunneled from the subcutaneous pocket to the most proximal muscle. There, a single recording channel and its reference wire were trimmed to length and stripped of insulation. The bare electrodes were inserted into the muscle belly perpendicular to the muscle striations, through 1-2cm of muscle. The electrodes were secured by suturing to the muscle at the insertion and exit points of the muscle belly. The remaining two channels were tunneled to the next muscle, where the process was repeated before the final electrode was tunneled to the most distal muscle and secured in the same manner. Finally, the telemetry unit was placed in the subcutaneous pocket on the upper rear flank and secured using a suture. Approximately 0.5g of Vancomycin powder was irrigated into the subcutaneous pocket, and the pocket was closed. The process was then repeated on the other side. After brief intraoperative testing, the animal was allowed to recover from anesthesia then returned to its pen.

#### Recording of EES-evoked motor potentials

At the beginning of the experimental session, the awake sheep was hoisted in a sling (Panepinto, Fort Collins, CO) until clearance between its hooves and the floor was observed. The HD64 was connected, enabling simultaneous stimulation, recording, and control of the multiplexer. Stimulation amplitude ranges were identified for each sheep, ranging from below motor threshold to the maximum comfortable response above motor threshold. Five stimulation amplitudes were selected in the comfortable range, and each stimulation was delivered at four frequencies (10, 25, 50, and 100 Hz).

In monopolar stimulation trials, stimulation was independently applied to each of the 60 contacts. A cathode-leading, 3:1 asymmetrical, charge-balanced waveform was used in all cases. The cathodic-phase pulse width was 167 μs, and the stimulation train duration was 300 ms. The stimulation anode was the implanted reference wire in the paraspinal muscles. The inter-train interval was randomly drawn from a uniform distribution spanning 1-2 seconds, and stimulus presentation order was randomized.

In bipolar stimulation trials, bipolar pairs were defined such that the spacing between the cathode and anode represented the minimum spacing possible on either the HD64 or Medtronic 5-6-5 paddles. Here, stimulation was provided at 5 amplitude values in the comfortable range for each sheep at a frequency of 10 Hz. This 100 ms inter-pulse-interval was selected to maximize the latency between stimuli so that the neural response to the previous pulse had subsided prior to the delivery of the subsequent stimulation. The stimulation waveform shape, train duration, and inter-train interval were unchanged from the monopolar trials. Stimulation presentation was again randomized.

Data was collected during stimulation across 60 electrode contacts per spinal paddle and 6-8 EMG channels. The spinal, EMG, and stimulation data were synchronized by injecting a known bitstream into the time series data of the spinal and muscular recordings. Stimulation sessions did not exceed three hours, and the sheep were constantly monitored for signs of distress and fed throughout each session. At the conclusion of recording, the animal was returned to their pen.

#### Recording of EES-evoked spinal potentials

At the beginning of each session, the awake sheep was hoisted in a sling, and the HD64 was connected as described previously. The HD64 was routed to create a stimulating bipole between the caudal-most midline electrodes and a recording bipole, either 7.3 mm, 17.0 mm, 26.7 mm, or 31.6 mm more rostral than the stimulating bipole. As the recording device (A-M Systems Model 1800) consisted of only two recording channels, each of the bipoles was recorded sequentially. For each stimulation event, a cathode-leading, symmetrical EES pulse was delivered to the stimulation bipole. The EES amplitude was set to 2 mA, 4 mA, and 6 mA, while the width of the cathodic phase was kept constant at 25 μs.

#### Recording of TENS-evoked spinal potentials

With the sheep hoisted in the sling apparatus, the bony anatomy of the right-side hind fetlock was palpated, and a ring of wool just proximal to the fetlock was shorn, extending approximately 10 cm proximally. The exposed skin was cleaned with isopropyl alcohol, which was allowed to air dry. 2”x 2” TENS (Transcutaneous Electrical Nerve Stimulation) patches (Balego, Minneapolis, MN) were cut to size, placed on the medial and lateral aspects of the cannon bone, then connected to the stimulator device (Model 4100, A-M Systems Inc, Carlsborg, WA). The stimulation monitoring channel was connected to an analog input on the EMG system for synchronization. Stimulation amplitude was set to 25V, and stimulation pulse width was 250 μs. The interstimulus interval was 500 ms. The above process was then repeated for the left hindlimb.

### Extension of state-of-the-art EES parameter inference neural networks

#### Model reparameterization and data acquisition

To extend the state-of-the-art EES parameter inference neural network models to 60 electrodes, the input space was reparameterized. The original model accepted a 3D input feature vector: amplitude, frequency, and electrode index (Govindarajan et al. 2022). The new model used here now accepts a 4D input vector: amplitude, frequency, electrode mediolateral coordinate, and electrode rostrocaudal coordinate (i.e., inputting an x-y location rather than hardcoding an arbitrary electrode number). By defining the electrode position on a continual space, positional relationships between each electrode could be learned. As in the original model, frequency and amplitude values were normalized to the range [0, 1).

For each sheep, radiographs were collected following the implantation of the HD64 electrodes. By examining the orientation of each electrode contact, an affine transformation was computed to account for skew between the electrode paddles and the radiograph camera. In this transformed space, the relative position of the rostral paddle (if placed) was determined with respect to the caudal paddle. Using the known dimensions of the HD64, the coordinates of each electrode contact were calculated. The electrode coordinates were normalized to the same range as the amplitude and frequency (the left-most caudal electrode contact had a coordinate of (0, 0), with increasing values extending rightward and rostrally).

The data acquisition phase proceeded as described for recording of monopolar EES-evoked motor responses.

#### Data handling & evaluation

The data handling and preprocessing steps were performed as previously described (Govindarajan et al. 2022). Briefly, the EMG recordings from bilateral *extensor digitorum longus*, *biceps femoris*, and *gastrocnemius* were high-pass filtered at 3 Hz, then a second-order infinite impulse response notch filter with a quality factor of 35 at 60 Hz. EMG envelopes were calculated by computing the moving RMS (root mean squared) value of each signal for a 300 ms window (50% window overlap). The enveloped data was epoched from 100 ms prior to the onset of stimulation to 300 ms after the conclusion of the stimulation train (700 ms epochs). The epoched data was labeled with the stimulation amplitude, frequency, and electrode coordinates determined by the map described previously. This dataset was the 100% density dataset, as it contained every stimulation electrode. The 50% density dataset was created by removing 30 electrode contacts from the 100% density dataset. Finally, the 25% density dataset was created by removing an additional 14 contacts from the 50% density dataset. The remaining data handling steps (outlier removal, unreliable sample removal, subthreshold EMG removal, and EMG summarization) were performed as previously described (Govindarajan et al. 2022). Independent models were then trained for each density dataset.

Following the completion of training and inference, stimulation proposals were generated for a set of target EMG responses obtained from testing EES parameter sets (40% of all conditions were held out, as in (Govindarajan et al. 2022)). Since these testing data were held out from the training data (the remaining 60% EES parameter conditions), the combination of target EMG responses and their EES parameters has never been shown to the network during the training phase. Proposals to achieve these target EMG responses were generated independently by all three models. If the stimulation contact coordinate inferred for a proposal was not encircled by a stimulation contact, the proposal was snapped to the nearest contact. The proposed stimulation amplitudes were checked to comply with the ranges the researcher found comfortable for each sheep and through software limits. To assess changes in the distribution of evoked motor responses, a set of stimulation parameters applied during the collection phase were repeated at the start of the evaluation phase. Then, each proposal was delivered 10 times, and the proposal order was randomized. After collection, the L1 error between the proposed EMG response and the achieved EMG response was calculated.

### Statistics and Data Processing

Following the completion of an experimental session, the recorded data were analyzed offline using custom-written code in MATLAB (R2023b) and Python 3.8.17 (using SciKit-Learn version 1.4.1 (Pedregosa et al. 2011) and Uniform Manifold Approximation and Projection (UMAP) module (McInnes et al. 2018)). To compute recruitment curves, the recorded EMG signals were split into 500 ms stimulation-triggered epochs. This epoch length was selected to capture the 300 ms pulse train and residual motor effects. The data was then rectified, and the power was recruited by taking the absolute value and then summing all samples in each epoch. The power values were then normalized to a range of [0, 1] for each muscle by subtracting the minimum activation and dividing by the range of activations. The EMG power (rectified area under the curve, or rAUC) was averaged across the four stimulation trials for each amplitude, frequency, and electrode combination, and the standard deviation was computed. To identify statistical clusters of evoked EMG responses, the single-trial recruitment data was rearranged into a 2D table, where rows were stimulation electrodes, and the recruitment data for each stimulation frequency was horizontally concatenated to form the columns. Raw rAUCs were normalized using the Yeo-Johnson power transformation (Yeo and Johnson 2000), and used as input features for the Uniform Manifold Approximation and Projection (UMAP) algorithm (McInnes, Healy, and Melville 2018; Gorman 2018). A two-dimensional embedding was generated, where each point represents the rAUC response pattern across muscles, stimulation amplitudes, and frequencies for a given electrode (four points per electrode, corresponding to the four stimulation repetitions). Spectral clustering (Shi and Malik 2000) was used to identify between 1 and 16 clusters of electrodes in the dataset, representing groups of electrodes with similar rAUC response patterns. A silhouette analysis (Rousseeuw 1987) indicated that the optimal number of clusters was eight. If the four repeats of the same electrode did not fall into the same cluster, that electrode was assigned the cluster label corresponding to the majority of the four labels. The process of epoching, rectifying. and integrating the EMG signal was repeated for the bipolar stimulation datasets. Before normalizing the bipolar data, the unnormalized monopolar data was included such that both monopolar and bipolar EMG responses were normalized to the same range. The EMG distributions for each stimulation configuration were compared using a Mann-Whitney U test. This non-parametric test was selected as it does not assume normality of the underlying distributions. Finally, the selectivity indexes were computed for each muscle in all stimulation electrode configurations (Badi et al. 2021; Bryson et al. 2023). The distributions of selectivity indexes were also compared using a Mann-Whitney U test.

Spinal evoked compound action potentials (ECAPs) and SEPs were first split into 500 μs and 100 ms stimulation-triggered epochs, respectively. The epochs were averaged across 50 stimulation trials and filtered with a low-pass filter at 2000 Hz. The mean and standard deviation of the ECAPs were calculated for each amplitude and recording bipole. The latency of the peak of the response was identified for each of the 50 single trials. A linear fit of the latency-distance relationship was calculated, and the gradient of this fit was calculated to determine the conduction velocity. The SEPs were re-referenced by subtracting the inverting electrode recording from the non-inverting electrode recording for each recording bipole. The mean SEP response was calculated for each recording channel. To assess the spatial correlation between recorded channels, the normalized zero-lag cross-correlation coefficient was computed between each recording channel for single trials. For each channel, the mean pairwise correlation to other channels was calculated over each trial. To assess the uniqueness between channels, the absolute value of the correlation was computed and then subtracted from 1. That is, perfectly correlated channels, with a mean pairwise correlation coefficient of 1, would receive a score of 0, while completely uncorrelated channels would receive a score of 1. The distribution of uniqueness scores was compared using a Kruskal-Wallis test. This non-parametric test was selected as it does not assume normality of the underlying distributions.

To evaluate the performance of the EMG response simulation neural network (the forward model), the L1 loss was computed by subtracting the predicted EMG vector from the evoked EMG vector and then taking the absolute value. This process occurred on a held-out dataset of mean EMG responses to 480 stimulation combinations, each with 4 repeats. The distributions were compared using a Kruskal-Wallis test. The random performance of each network was determined by generating a randomized prediction for each muscle from a uniform distribution ranging from 0 to the maximum response in the training dataset. Then, this randomized EMG vector was compared to the target vector. The L1 losses were split for each model by inclusion of the stimulating electrode in the training dataset. The L1 losses were compared within models using the Mann-Whittney U test. Finally, the L1 losses were split for each model by muscle, and their distributions were compared using a Kruskal-Wallis test. To evaluate the performance of the parameter inference neural network (the inverse model), the rAUC was computed for each of the responses to inferred parameters. The mean L1 error between the predicted and achieved EMG rAUC vectors was calculated across muscles. Additionally, the L1 error between the EMG responses to the original EES parameters and the responses to these same parameters sent during the evaluation session (which occurred some hours after the training session) was calculated. The distributions of L1 errors were compared using the Kruskal-Wallis test.

## Results

### Successful chronic implantation of the HD64

To evaluate the safety of chronic implantation of the HD64 devices, we monitored the condition of the sheep continuously throughout the study and evaluated our ability to control the multiplexer on the active array. We implanted two sheep with 1 (S1) or 2 (S2) HD64 EES paddles which remained implanted for 15 months. The multiplexers were configured 1598 and 4562 times on 34 and 51 unique days, respectively, throughout the study. No device-related malfunctions were observed, and control of the multiplexer remained stable. The impedance from the distal connector end of the lead-tails to the contacts (including the paths through the multiplexer) were measured at several points up to 274 days post implant. Over this time, no significant changes in the distribution of impedances were observed (*p* > 0.05, Mann-Whitney U test) (Fig. 2d). After this time, we changed stimulator devices, and the second device did not have impedance measurement capabilities. While the HD64 was implanted in the animals, we observed consistent stimulation to EMG responses and spinal responses to spinal and peripheral stimulation.

### Active multiplexing enables precise motor recruitment

After implantation, we evaluated the benefit of a high-density electrode paddle in generating targeted motor recruitment from EES by recording motor-evoked responses from 6-8 bilateral lower extremity EMG sensors. Following EES, the EMG signal was rectified, then the area under the curve (rAUC) was calculated and plotted as a function of amplitude (Fig. 3a). Applying monopolar EES to each of the 60 contacts in S1 produced a detectable motor response in at least 1 of 6 muscles. For each electrode contact, the EES amplitude required to reach 33% of the maximum activation for each muscle was identified (Fig. 3b). Contacts marked N.R. could not recruit the muscle to 33% of its maximum activation. Our analysis identified a lower EES amplitude required to recruit ipsilateral muscles than contralateral muscles, consistent with previous literature (Calvert et al. 2019). The results for S2 are presented in Supplemental Figure 1.

**Figure 3.**
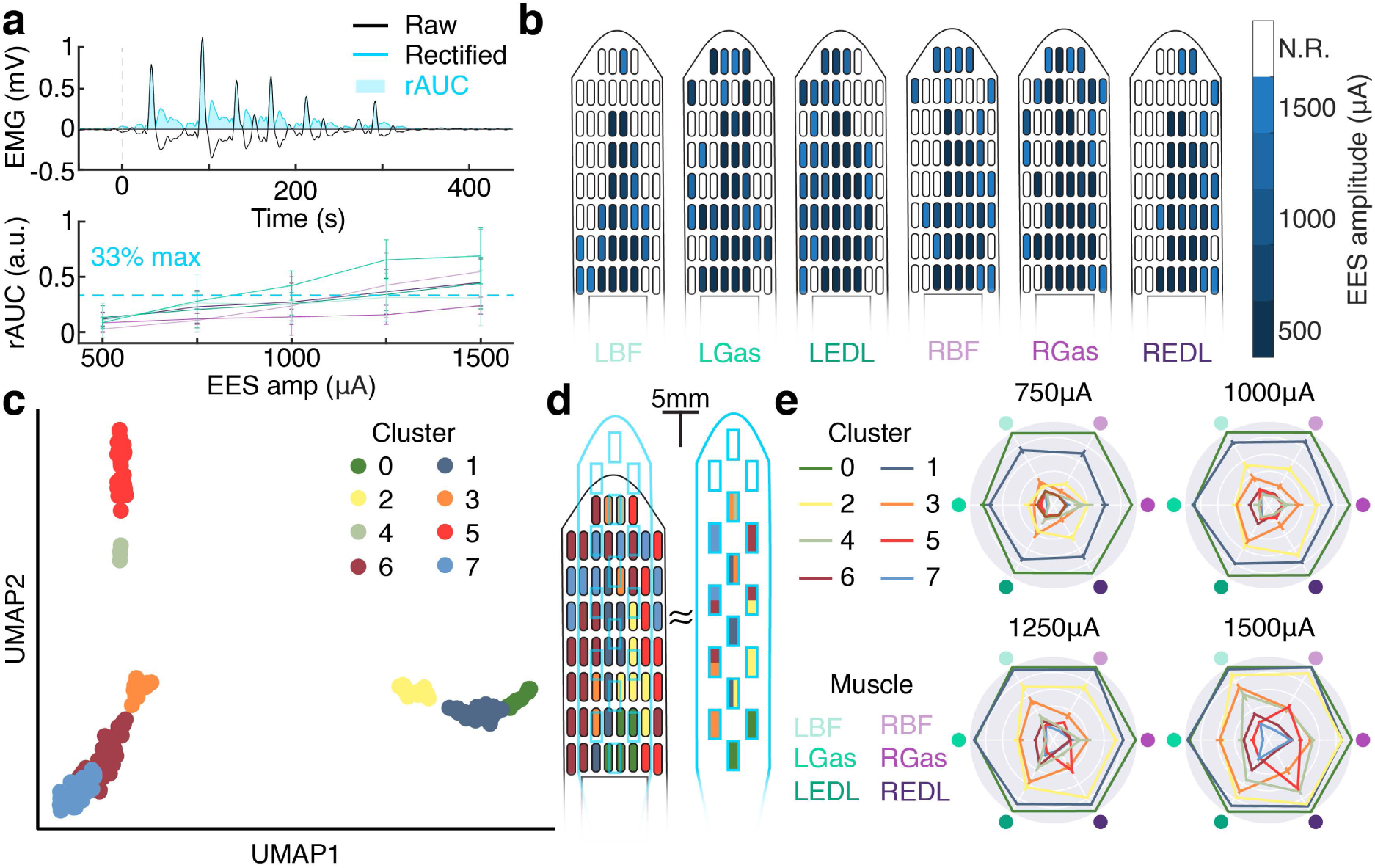
Clustering analysis of motor outputs evoked by monopolar stimulation. **a)** A graphical overview of the electromyography (EMG) preprocessing. A representative EMG trace (black) is rectified (blue trace), then the rectified area under the curve is calculated (blue shaded region). A sample recruitment curve is shown for six muscles. The points are the mean, and the bars show the standard deviation. A blue dashed line indicates the 33% of maximum activation threshold used for panel b. **b)** The minimum required epidural electrical stimulation (EES) amplitude required to recruit each of the six instrumented muscles to 33% of their maximum activation. Electrodes marked “N.R.” could not recruit the muscle to 33% of its maximum activation at the amplitude range tested. **c)** A 2D projection of the embedded recruitment data. Each point is a stimulation event, and the points are colored by which cluster the stimulating electrode resides in. **d)** A diagram of the HD64, with the electrode contacts colored by which cluster the electrode resides in. A scale drawing of the Medtronic 5-6-5 is overlaid to highlight which cluster(s) each of the 5-6-5’s electrodes contact. **e)** Representative radar plots (EES frequency = 50Hz) show the mean responses for each stimulation contact cluster. The clusters exhibit diverse recruitment patterns and relationships with amplitude.

The spatial diversity of EES-evoked motor responses was examined by performing unsupervised clustering. Uniform Manifold Approximation and Projection (UMAP) dimensionality reduction was performed on the recruitment data for each of the 60 electrodes. The low-dimensional embeddings were then clustered by statistical similarity. Eight independent clusters were identified. Electrodes for each cluster exhibited colocalization (Fig. 3c, Supplemental Figure 2) on the HD64 paddle. To simulate the performance of a commercially available EES paddle (5-6-5, Medtronic, Minneapolis, MN), a scale drawing of the commercial paddle was overlaid on the HD64 clusters. While some commercial paddle contacts overlaid a single cluster, many contacts straddled two or more independent clusters, and Cluster 5 was not reachable (Fig. 3d). Consequently, the motor responses evoked using the commercial paddle may lack the fidelity available using the HD64. To explore the diversity of the recruitment patterns across clusters, the mean recruitment for each electrode in each cluster was calculated and then normalized using the Yeo-Johnson power transformation (Yeo and Johnson 2000). Fig. 3e provides an example of the recruitment patterns observed for each cluster at a stimulation frequency of 50 Hz.

To further explore the stimulation fidelity afforded using a high-density electrode paddle compared to a commercially available paddle, we examined the effect of current steering. Our stimulating analog front-ends were equipped with a single current source, enabling us to create a single bipole. When no anodic contact was specified, current returned through the ground wire implanted in the paraspinal muscles. This creates a much more dispersed electric field than if the current return was more proximal to the cathodic contact. By using an anodic stimulation contact, current is simultaneously sourced and sinked from the cathodic and anodic contacts, respectively. Thus, the electric field is “steered” towards the anodic contact, changing the intensity and dispersion of the electric field. EES was delivered using bipolar stimulation pairs spaced using the minimum inter-electrode spacing available on the HD64 (“narrow”, Supp Fig. 3a and 3g), or the minimum spacing available on the 5-6-5 (“wide”, Supp Fig. 3d and 3g), and was compared to each other, or a monopolar baseline (Supp Fig. 3a and 3d). For each lower extremity muscle, the rAUC and selectivity of the response to stimulation was calculated, as described previously. Using a Mann-Whitney U test, a significant difference was identified between the narrow and wide bipolar stimulations for all stimulation amplitudes (*p* < 0.05), though never in all muscles, indicative of a difference in the selectivity of activation (Supp Fig. 3h). Comparing the selectivity of the narrow and wide bipolar fields, a significant difference was identified in at least one muscle for all stimulation amplitudes. At the maximum EES amplitude tested (1500 μA) 66% of instrumented muscles demonstrated significantly different response selectivity (Supp Fig. 3i). Other bipolar arrangements are presented in Supplemental Figure 4.

### Active multiplexing enables localized referencing

To evaluate the benefits of dense recording bipoles, we began examining epidural spinal responses to spinal and peripheral stimulation. The experimental configuration for this section is shown in Fig. 4a. To examine the propagation of evoked compound action potentials (ECAPs) on the spinal cord using the HD64, a EES bipole was created on the caudal midline of the paddle. Four bipolar recording pairs were established along the midline of the paddle, with centers distanced 7.3 mm, 17.0 mm, 26.7 mm, and 31.6 mm rostrally from the center of the stimulating bipole respectively (Fig. 4b). Single cathode-leading, biphasic, symmetrical EES pulses were delivered with amplitudes 2 mA, 4 mA, and 6 mA. The pulse width remained constant at 25 μs. The responses recorded by each bipole are presented in Fig. 4c. The elapsed time between stimulation and the peak of the ECAP was identified at each bipole for each of the 50 trials (Fig. 4d). Using the distance between the stimulating and recording bipoles and the time of peak response, the conduction velocity of the evoked response was calculated. For stimulation amplitudes 2 mA, 4 mA, and 6 mA, the conduction velocities were 114.9 m/s, 114.6 m/s, and 111.8 m/s, respectively.

**Figure 4.**
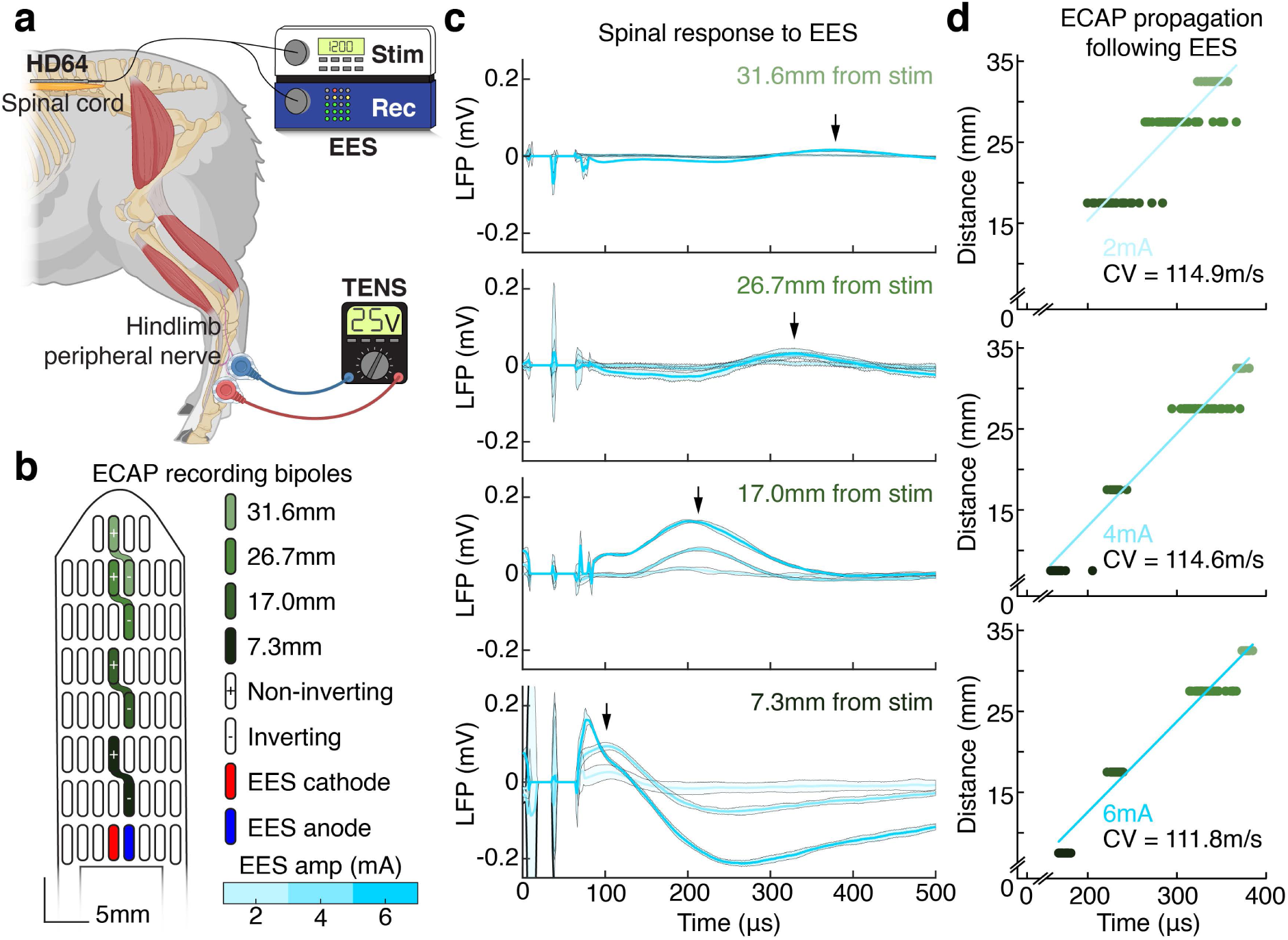
Assessing spinal responses detected by monopolar, wide bipolar, and narrow bipolar recordings. **a)** A graphical overview of the spinal local field potential (LFP) recording, epidural electrical stimulation (EES) and transcutaneous electrical nerve stimulation (TENS) setup. **b)** A map of the HD64, showing the stimulation cathode (red) and anode (blue). The recording bipoles and their distances from stimulation are shown. **c)** The mean and standard deviation of the spinal evoked compound action potential (ECAP) response recorded using various bipolar pairs along the midline of the spinal cord, for three stimulation amplitudes. The peak of the 4mA stimulation response is shown with a black arrow. Note that peak latency increases with separation from the stimulation bipole. **d)** The peak latencies of 50 single trials, plotted as a function of separation from the stimulating bipole, for each stimulation amplitude. The samples are colored by their recording bipole. A linear fit is plotted in blue, with the conduction velocity (CV) inset.

We examined the SEPs in response to peripheral application of TENS using three recording strategies, each containing six recording channels (Supp. Fig. 5a). The monopolar recording strategy used six consecutive HD64 contacts, with a recording reference wire implanted in the epidural space. The bipolar HD64 arrangement created six consecutive bipolar recording pairs using the minimum inter-electrode spacing available on the HD64. Finally, the bipolar 5-6-5 arrangement simulated the best-case bipolar arrangement using a commercial spinal paddle by creating six bipolar recording pairs with an interelectrode distance similar to that found on the Medtronic 5-6-5 (a to-scale drawing of this paddle is presented in blue in Supp. Fig. 5a). Supplementary Fig. 5b shows the trial average (50 trials) SEP in response to 25V TENS applied to the right fetlock for all three arrangements. While the SEP is clearly prominent in the monopolar arrangement, the responses recorded by all channels are highly correlated. To identify the degree of correlation in recorded SEPs between channels, the mean pairwise zero-lag cross-correlation coefficient was calculated for each channel on single trials. The effect of arbitrary bipole orientation was removed by taking the absolute value of the correlation coefficient. A statistically significant difference between the distribution of correlations was identified for all recording arrangements (*p* < 0.05, Kruskal–Wallis test) (Supp. Fig. 5c). Spectral comparisons are made in Supplemental Figure 6.

### Continuous encoding of electrode position enables inference over electrode space

Increasing the number of stimulation contacts increases the solution space of automated parameter inference neural networks. We leveraged the dense grid of contacts on the HD64 to enable spatial encoding of EES parameters during training and inference. The reparameterized neural network models were successfully trained to infer evoked EMG power following EES for all three electrode densities (Fig. 5a). The cumulative L1 loss between the predicted and actual EMG power for the held-out dataset is shown at the top of Fig. 5b for each model. Significant differences were found between the 25% model and all other models (*p* < 0.05, Kruskal–Wallis test), but not between the 50% and 100% models (*p* > 0.05, Kruskal–Wallis test). Examining the L1 prediction error for each model, a significantly higher error was produced when predicting responses to stimulation on unseen electrodes for the 25% model (*p* < 0.05, Mann-Whitney U test). This was not present for the 50% model, indicating the 50% model’s ability to accurately infer EMG responses for unseen stimulation sites (*p* > 0.05, Mann-Whitney U test) (Fig. 5b bottom). In all cases, the prediction error was less than random chance (indicated by blue dots in Fig. 5b). Decomposing the L1 loss by muscle reveals the same result identified previously, where the only significant difference in L1 distribution occurs between the 25% model and the remaining models (*p* < 0.05, Kruskal–Wallis test), except for the left gastrocnemius, where prediction loss followed the same distribution for all models (*p* > 0.05, Kruskal–Wallis test) (Supp. Fig. 7).

**Figure 5.**
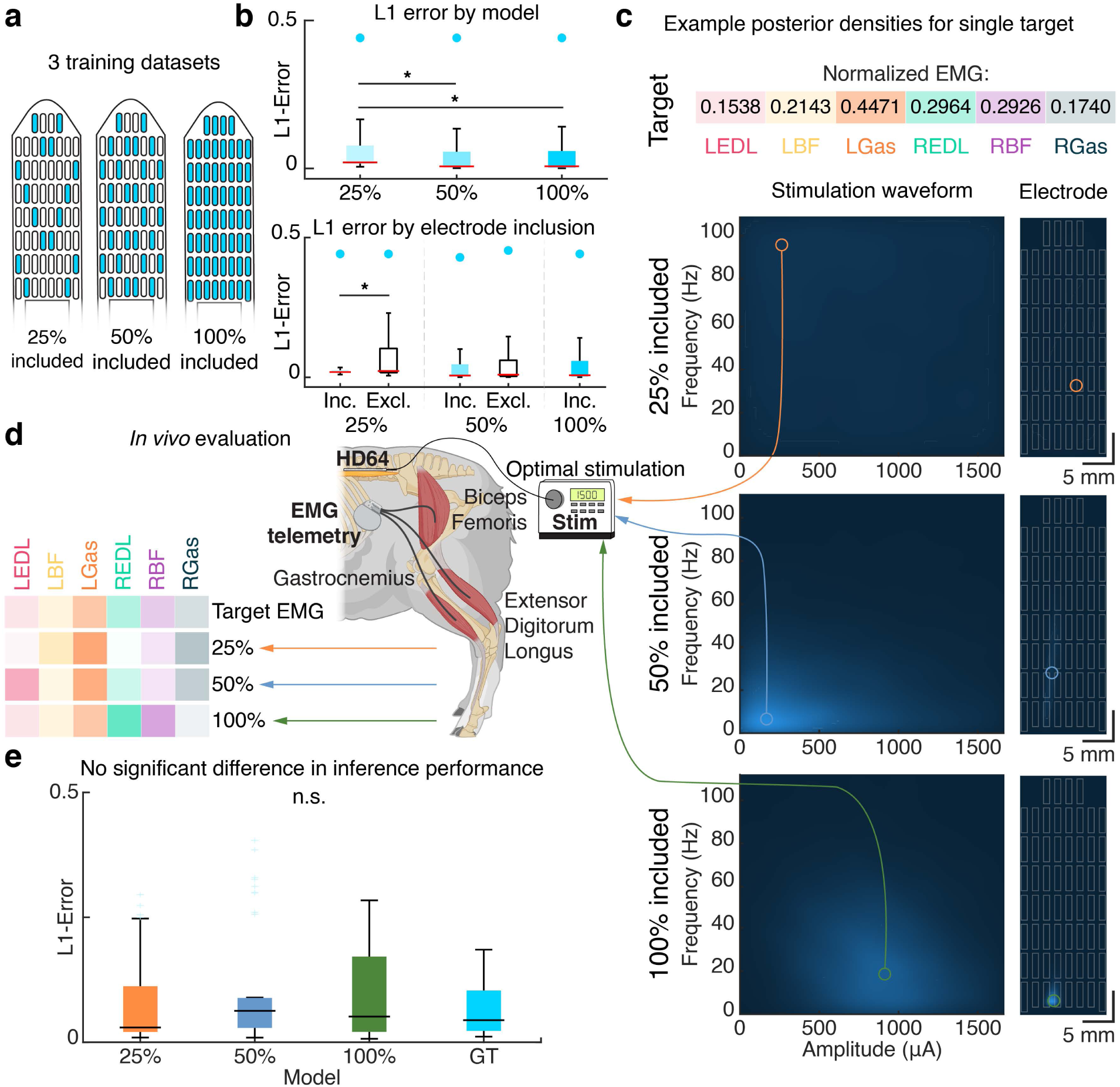
Evaluating the performance of the forward neural network model to predict sensorimotor computations. In all boxplots the horizontal bar is the median, the boxes extend from the 25th to the 75th percentile, and the blue dots represent a “random” baseline performance. Statistical significance was calculated using a Kruskal-Wallis non-parametric test, with a maximum *p*-value of 0.05. **a)** A map of the electrode contacts included in the training dataset for each model. Included electrodes are shaded blue. **b)** (top) The distribution of L1 errors between the ground truth and simulated electromyography (EMG) for a held out dataset of 480 stimulation parameter combinations for the three models. (bottom) The data in (top) split by inclusion of the stimulating electrode in the training dataset. **c)** An example target normalized electromyography (EMG) vector is shown at the top. The posterior densities over are shown for 3 models. Higher likelihood regions are indicated in lighter colors. **d)** The *in vivo* evaluation procedure. The posterior densities of each model are sampled to produce proposed stimulation parameter combinations for each target. These parameters are applied *in vivo*, and the evoked EMG vector is calculated. The L1-error between the target EMG vector and the evoked EMG vector is then computed. **e)** The distribution of mean L1 errors between the target EMG vector and the evoked EMG vector for each of the three models, and between EMG vectors evoked using identical stimuli in the training and evaluation sessions (“ground truth”, or GT).

Following successful amortization of spinal sensorimotor computations by the forward models, we performed inference using the inverse models to identify EES parameters predicted to evoke target EMG responses. The inverse models produced joint posterior density functions over stimulation parameters and electrode spaces for 32 target EMG vectors. Fig. 5c compares the posterior densities across the three models for a single EMG target. Posteriors for additional EMG targets are provided in Supplemental Figure 8. Here, the 25% model predicts low likelihood across all electrode-and parameter-space. As the number of included electrode positions increases, the model localizes an optimal stimulation location, indicated by the focal region of high likelihood. To evaluate the performance of the models, we selected target EMG vectors from the held-out dataset for evaluation *in vivo*. The workflow and representative results are outlined in Fig. 5d. The stimulation parameter and electrode combination that produced the highest likelihood is identified and then delivered using the stimulator device. The evoked EMG responses from the six instrumented muscles are recorded and then compared offline to the target EMG vector. This process was repeated for all three models. We then compared the L1 error between the target EMG response and the evoked EMG response across models (Fig. 5e). A nonparametric Kruskal-Wallis test revealed that the underlying distributions of L1 errors between targets and the responses were not significantly different between models (*p* > 0.05). This indicates that adequate training data to infer stimulation parameters to faithfully reproduce target EMG activity could be collected using a sparse subset (25%) of the available stimulation contacts. Further, no significant differences were observed between the re-sent stimulation parameter (the “ground truth”) distribution and the proposal distributions. This indicates that the parameter inference performance approached the soft upper bound given by the natural variations in EMG recorded across time.

## Discussion

In this work, we presented a novel, high-density, active electronic epidural electrical stimulation paddle, the HD64. We evaluated the hermeticity of the active electronics assembly, and found leak rates 15 times smaller than the threshold defined by MIL-STD-883K. Following ISO 10993-1:2018 biocompatibility testing, we confidently progressed to *in vivo* testing. Using the onboard electronics, we quantified the improvements in the diversity and selectivity of motor responses evoked using the HD64 compared to wider bipolar configurations. Further, we demonstrated the capabilities of the HD64 to provide superior referencing of somatosensory evoked potentials following peripheral nerve stimulation, significantly reducing the mean spatial correlation compared to bipoles with similar spacing to commercial paddles. Finally, we extended the current state-of-the-art stimulation parameter inference model to encode sensorimotor-EES relations, enabling increased spatial resolution of inferred EES parameters and utilizing 2.5x higher channel count compared to prior preclinical work and clinical standards.

Looking forward, our work here is a fundamental demonstration of active electronics embedded into neural interface electrodes themselves. While our work leverages multiplexing to achieve a high channel count, other architectures may use the same hermetic approach to integrate digital signal processing (DSP) or stimulation control directly into electrodes. We expect others to continue the path towards highly functional integrated neural interfaces to restore lost function following trauma or disease.

### Active electronics can be chronically hermetically sealed in an EES paddle

Our novel paddle contains an active multiplexing chip hermetically sealed on the body of the EES paddle, enabling communication with 60 bidirectional contacts through two 12-contact percutaneous lead tails. The increased contact density combinatoriality increases the number of possible stimulation configurations available for research and therapeutic applications. In contrast to this reconfigurability, current clinically available EES paddles sacrifice spatial resolution in favor of a general, “one-size-fits-most” approach. In research studies, prior work has suggested a library of paddles of different sizes to suit person-to-person variability in anatomy (Rowald et al. 2022). Instead, the reconfigurability of the HD64 allows for localized concentration of stimulating electrodes to activate a specific area of the spinal cord, enabling focal stimulation with current steering to prevent off-target effects without sacrificing the ability to simultaneously stimulate other distal regions.

As the first example of an EES paddle including active electronics, we were particularly interested in assessing the ability of the encapsulant to maintain patency during chronic implantation. We found that the HD64 remained functional for at least 15 months after implantation and did not display symptoms of fluid ingress. Communication between the externalized stimulation and recording hardware and the onboard multiplexer remained robust throughout the study. This initial result is promising for the future of epidural medical devices containing active electronics. With current technologies, chips of similar dimensions to the present device are capable of significant computational power. Our results may open the door for diverse advances in on-paddle processing, including signal processing or closed-loop stimulation parameter control, as has been hypothesized in prior literature (Zhang, Savolainen, and Constandinou 2022). While additional testing and regulatory approval is required before such technologies can be translated into clinical applications, this demonstration represents a promising first step.

### Onboard active multiplexing enables chronic stimulation-evoked responses

Our study benefits from decades of prior research showing the selective activation of motor neuron pools using EES applied to the lumbosacral spinal cord. Specifically, EES has been utilized heavily to restore walking ability following a spinal cord injury (Angeli et al. 2018; Capogrosso et al. 2016; Gill et al. 2018). Other studies have used EES to establish somatotopic maps of stimulation location to recruit lower limb extremities (Hofstoetter et al. 2021). Extending the mediolateral span through which EES can be delivered, while maintaining a high contact-density, enabled us to specifically target lateral structures of the spinal cord. Stimulation at these lateral sites generated stereotyped EMG responses statistically different from those achieved by medial stimulation, as determined by our spectral clustering analysis. These differences highlight the importance of lateral stimulation sites for enhancing the capability of locomotor-restorative EES. Such sites are not present on stimulation paddle arrays designed for chronic pain management.

Recording EES-evoked spinal responses, or ECAPs, can provide quantitative data regarding EES paddle status. ECAPs have been used as a control signal for pain-modulating EES (Mekhail et al. 2020), and we have previously suggested their utility in the identification of neural anatomy (Calvert et al. 2023). Using bipolar recording configurations on the HD64, we observed an orthodromically propagating response to EES. The conduction velocities of these responses are consistent with the published ranges for Type Ia and Ib axons, which have been shown to be recruited by EES in computational models (Capogrosso et al. 2013; Parker et al. 2020). Further, our work demonstrates the formation of dense recording bipoles on the ovine spinal cord, and uses these recording bipoles to examine EES- and TENS-evoked spinal responses. This is timely for the field of neural interfaces, with the recent advent of high-density neural probes (Jun et al. 2017; Steinmetz et al. 2021; Chamanzar et al. 2023) and electrocorticography (ECoG) grids (Rachinskiy et al. 2022; Palopoli-Trojani et al. 2024). Using conventional EES paddles, contacts are sparsely distributed to ensure coverage of a large area using a small number of electrodes. This sets a high minimum limit on contact separation.

When considering epidural electrical stimulation or sensing, we suggest that the active multiplexing on the HD64 enables dense bipoles to be created at arbitrary points of interest. Importantly, contact separation can be non-uniform (for example, tight spacing in regions of high spatial variability and wider spacing in less volatile regions) or change with time or desired function. These “activity-dependent” paddle arrangements allow for the capabilities of the stimulation system to be adjusted to suit the needs of patients when completing various activities of daily living (for example, greater trunk control during sitting vs. lower extremity motor control during locomotion in patients with motor dysfunction).

### Spatial encoding of EES enables parameter inference over arbitrary channels

Stimulation parameter inference remains an obstacle to widespread clinical implementation of EES (Solinsky, Specker-Sullivan, and Wexler 2020). Our previous parameter selection automation work (Govindarajan et al. 2022), and other subsequent literature (Bonizzato et al. 2023) treat stimulating electrodes as discrete variables. As such, spatial relationships (such as those uncovered in Fig. 3d) cannot be leveraged by the model. In this work, we instead evaluated a novel electrode encoding scheme that accounts for spatial relationships between electrode contacts. This approach is extensible to very high channel counts and allows the network to identify optimal stimulation locations not included in the training dataset. Using the results shown here, active electronics conducting neural network inference may be hermetically sealed into the paddle array body, enabling stimulation parameter inference to be conducted without additional hardware or communication requirements.

As the number of available stimulation channels increases, the time required to collect training data from every contact scales linearly. Unaddressed, this is a barrier to clinical translation of parameter selection automation techniques and high-density EES paddles. Here, our work shows that exhaustive sampling of every stimulation contact is not required if the electrode encoding scheme is continuous. No significant decrease in EMG response prediction performance is observed for electrode inclusion rates as low as 50%, and stimulation parameter inference performance did not significantly vary across inclusion rates tested. The development of efficient machine learning tools, such as the one presented in this manuscript, greatly reduces the amount of training data required. Increasing efficiency and decreasing data collection efforts will aid in scaling EES for clinical use, which could yield meaningful benefits during spinal rehabilitation in patients with neural dysfunction.

### Study limitations and implications for future research

Our study introduces a high-density smart EES paddle with an integrated multiplexer and demonstrates the utility of increased electrode density. However, several limitations should be considered. While the characterization presented in this manuscript empirically supports the use of high-density electrode paddles in these applications, a more thorough assessment may be possible with the addition of computational models. Previous advances in electrode design were informed by finite element models of 15 healthy volunteers (Rowald et al. 2022). Such models could quantify recruitment of off-target motor neuron pools after applying stimulation at bipoles of various sizes, however the work presented here utilizes an ovine model. The spinal cord of the sheep is similar to that of humans in terms of gross anatomy and size and has been used as a model for the study of spinal electrophysiology (Parker et al. 2013, 2020; Chakravarthy, Bink, and Dinsmoor 2020; Calvert et al. 2023), but the results of this manuscript may not be generalizable to human anatomy. Future studies are required to evaluate the therapeutic potential of high-density EES electrodes or to quantify the effect of stimulation focality on behavioral values such as sensation. One current barrier to clinical translation of the HD64 is the lack of existing implantable pulse generators (IPGs) capable of generating the power and communication signals necessary to control the onboard multiplexer. Additionally, the inclusion of active electronics in implanted electrodes increases the power draw of the implantable system, and must be considered when selecting an appropriate IPG battery size, though the HD64 consumes only 20μA in steady-state operation and 100μA when reconfiguring connections. Our neural network models demonstrate sufficient accuracy to match EMG patterns, however, these experiments were conducted with the sheep at rest and elevated in a sling. A more functional task may instead consider applying a stimulation sequence to match a target kinematic trajectory. Additionally, our neural network parameter inference is limited to monopolar stimulation patterns and, therefore, cannot take advantage of the improved selectivity identified by bipolar stimulation in this manuscript. Bipolar stimulation dramatically increases the number of electrode configurations the network must evaluate; the additional complexity of which is a promising target for future research.

## Conclusions

In summary, we successfully designed and chronically implanted a high-density smart EES paddle, the HD64, which is the first to integrate active electronics into a hermetic package on the spinal cord. Our hermetic assembly was tested to exceed standards for active hermetic electronics. During chronic *in vivo* implantation, no device-related malfunctions, nor symptoms of moisture ingress were observed, enabling demonstration of fundamental stimulation-response characteristics in the ovine spinal cord. These results highlight the translational potential of high-density EES paddles, and provide a foundation for more advanced computation and processing to be integrated directly into neural interfaces.

## Acknowledgments

We would like to thank Aaron Gregoire and the Center for Animal Resources and Education (CARE) staff at Brown University for their assistance with animal care, as well as Alison Gorbatov and Krista Tocco from the Center for Neurorestoration and Neurotechnology (CfNN) for their assistance with the Providence VA Medical System regulatory work. We would also like to thank John Murphy for his engineering expertise in designing modifications to the sling apparatus. Finally, we would like to thank Moses Goddard for his surgical time and expertise in implanting the EMG system. Figs. 2a, 4ac, and 5d were created with BioRender.com.

## Data availability statement

The data that support the findings of this study are available from Zenodo at https://doi.org/10.5281/zenodo.11223073. Matlab scripts for reproducing the findings of this study are available at https://github.com/neuromotion/The_HD64.

## Authorship statement

SRP: Conceptualization, Methodology, Software, Formal analysis, Investigation, Data Curation, Writing - original draft, Visualization. JSC: Conceptualization, Methodology, Software, Formal analysis, Investigation, Data Curation, Writing - review & editing. RD: Conceptualization, Methodology, Software, Formal analysis, Investigation, Data Curation, Writing - review & editing. JJ: Conceptualization, Methodology, Software, Investigation, Writing - review & editing. LNG: Conceptualization, Methodology, Software, Investigation, Writing - review & editing. KA: Conceptualization, Methodology, Validation, Writing - review & editing. GC: Conceptualization, Methodology, Validation, Writing - review & editing. YI: Conceptualization, Methodology, Validation, Supervision, Writing - review & editing. ES: Conceptualization, Investigation, Writing - review & editing. JSF: Conceptualization, Investigation, Supervision, Funding acquisition, Writing - review & editing. TS: Conceptualization, Supervision, Funding acquisition, Writing - review & editing. DAB: Conceptualization, Methodology, Investigation, Resources, Writing - original draft, Writing - review & editing, Supervision, Project administration, Funding acquisition. BM: Conceptualization, Methodology, Investigation, Resources, Writing - original draft, Writing - review & editing, Supervision, Project administration, Funding acquisition.

DAB and BM contributed equally to this manuscript. Final approval was given by all authors.

## Supplementary Information

**Supplementary Table 1.** A summary of the biocompatibility tests and test results completed by a contracted test house. Tests were performed according to the surface area (specified by the ISO 10993 test method) and a specific number of electrodes were manufactured to satisfy the surface area requirement. In the case of pyrogenicity, for example, approximately 30 paddle electrodes were manufactured to perform the tests. Each ISO 10993 test specifies the test criteria and outcomes. The HD64 passed all tests as per ISO 10993.

| **Endpoints for Biological Evaluation** | **Reference** | **Test Method** | **Extraction Method** | **Result** |
| --- | --- | --- | --- | --- |
| Irritation | ISO 10993-10 | ISO Intracutaneous Reactivity Test, 120 cm<sup>2</sup> of devices | 50°C, 72 hrs, 3 cm<sup>2</sup>/mL<br>0.9% Sodium Chloride Solution, Sesame Oil, NF | PASS |
| Material-Mediated pyrogenicity | ISO 10993-11 | Material-Mediated Pyrogenicity, 270 cm<sup>2</sup> of devices | 50°C, 72 hrs, 3 cm<sup>2</sup>/mL Sterile Non-Pyrogenic Saline (SNPS) | PASS |
| Acute Systemic Toxicity | ISO 10993-11 | Acute Systemic Toxicity, 60 cm<sup>2</sup> of devices | 50°C, 72 hrs, 3 cm<sup>2</sup>/mL<br>0.9% Sodium Chloride Solution, Sesame Oil, NF | PASS |
| Implantation | ISO 10993-6 | Subcutaneous Implantation Study (30 days), 12+ device samples | Subcutaneous Implantation with Histopathology<br>2 sites/rabbit, HDPE control<br>2 sites/rabbit | PASS |
| Genotoxicity | ISO 10993-3 | Ames Assay, 60 cm<sup>2</sup> of devices | 50°C, 72 hrs, 3 cm<sup>2</sup>/mL, NS, PEG | PASS |
| Genotoxicity | ISO 10993-3 | Mouse Lymphoma Assay, 120 cm<sup>2</sup> of devices | 37°C, 72 hrs, 3 cm<sup>2</sup>/mL, Serum Free Cell Culture Medium, PEG | PASS |
| Ethylene Oxide Sterilization Residuals | ISO 10993-7 | HS GC/MS Limited exp: ETO<20 mg/day, | Assurance level of 10<sup>-6</sup> after 24 hours of heated aeration. Device presented | PASS |

| Endpoints for Biological Evaluation | Reference | Test Method | Extraction Method | Result |
| --- | --- | --- | --- | --- |
|  |  | ECH<12mg/day<br>Prolonged:<br><2mg/day,<br><20 mg in 24 hours.<br>ECH<2 mg/day,<br><12mg<br>in 24 hours, <60 mg in 30 days., 5+ devices | an equivalent or lesser challenge than an existing validated process challenge device. |  |
| Bacterial endotoxin level (BET) | USP | LAL Assay (<2.15 EU/device for potential CSF contact), 2+ devices per lot | 2 lots, 3 samples <2 EU/ml | PASS |

**Supplemental Figure 1.**
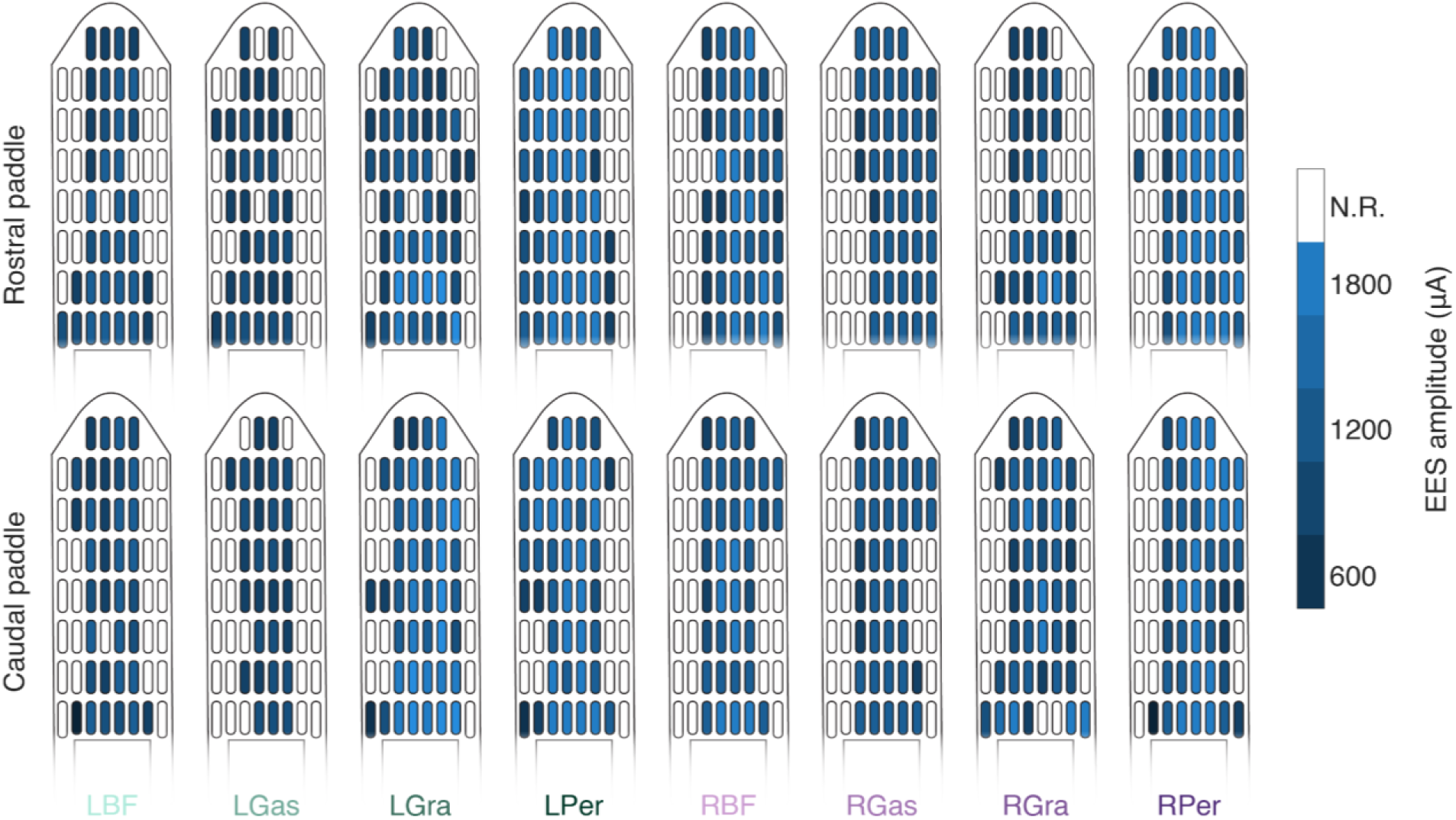
Epidural electrical stimulation amplitude thresholds for S2. Electrode contacts on the rostral and caudal paddles are colored by the minimum required epidural electrical stimulation (EES) amplitude required to recruit each of the six instrumented muscles to 33% of their maximum activation. Electrodes marked “N.R.” could not recruit the muscle to 33% of its maximum activation at the amplitude range tested.

**Supplemental Figure 2.**
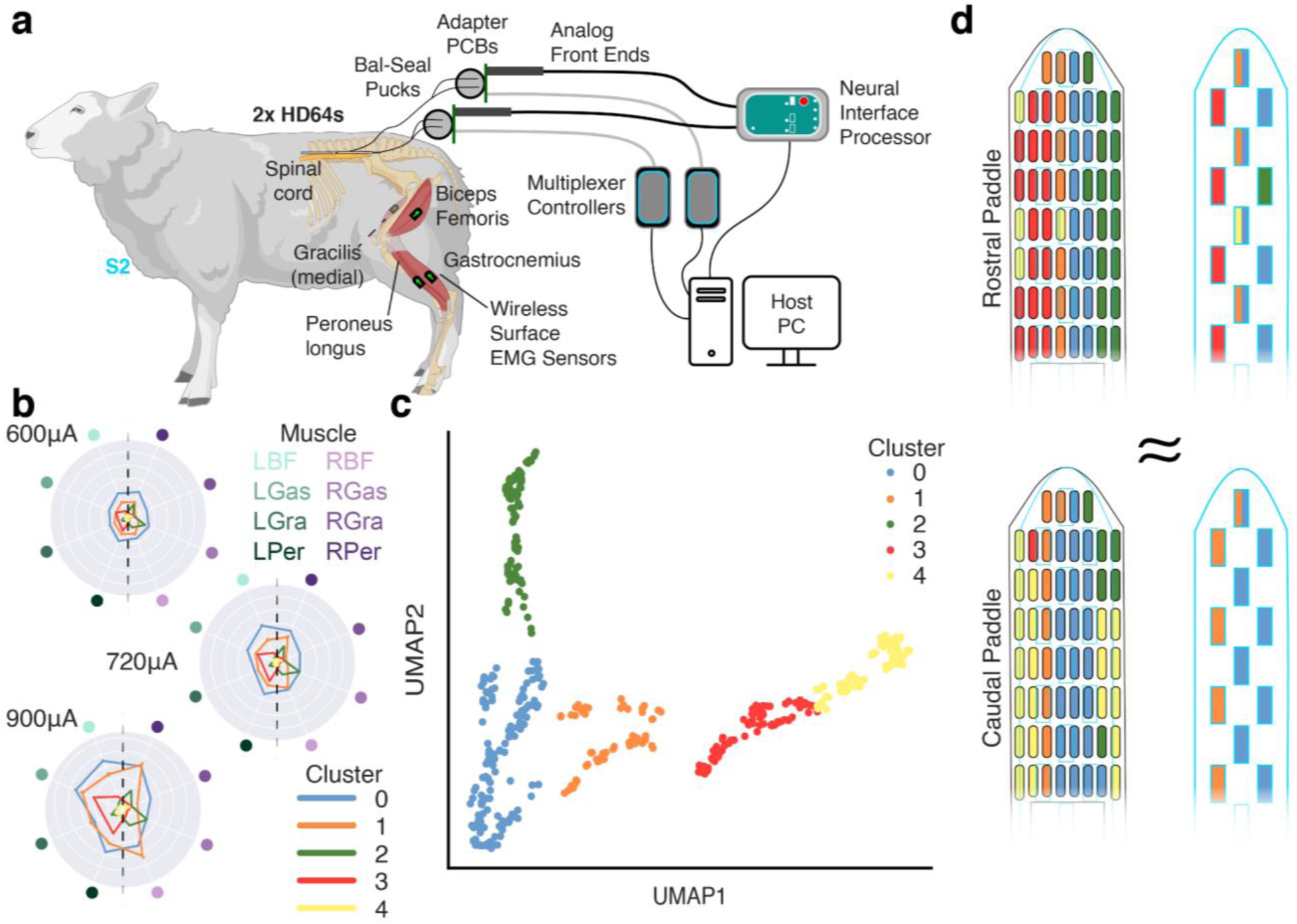
Algorithmic clustering of stimulation-evoked motor responses in S2. (a) A diagram of the experimental setup. In contrast to S1, S2 uses surface electromyography (EMG) sensors, instrumenting four bilateral hindlimb muscles, and stimulation is applied on two HD64 paddles. (b) Representative radar plots (EES frequency = 50Hz) show the mean responses for each stimulation contact cluster. The clusters exhibit diverse recruitment patterns and relationships with amplitude. (c) A 2D projection of the embedded recruitment data. Each point is a stimulation event, and the points are colored by which cluster the stimulating electrode resides in. (d) A diagram of the two HD64s, with the electrode contacts colored by which cluster the electrode resides in. A scale drawing of the Medtronic 5-6-5 is overlaid to highlight which cluster(s) each of the 5-6-5’s electrodes contact.

**Supplemental Figure 3.**
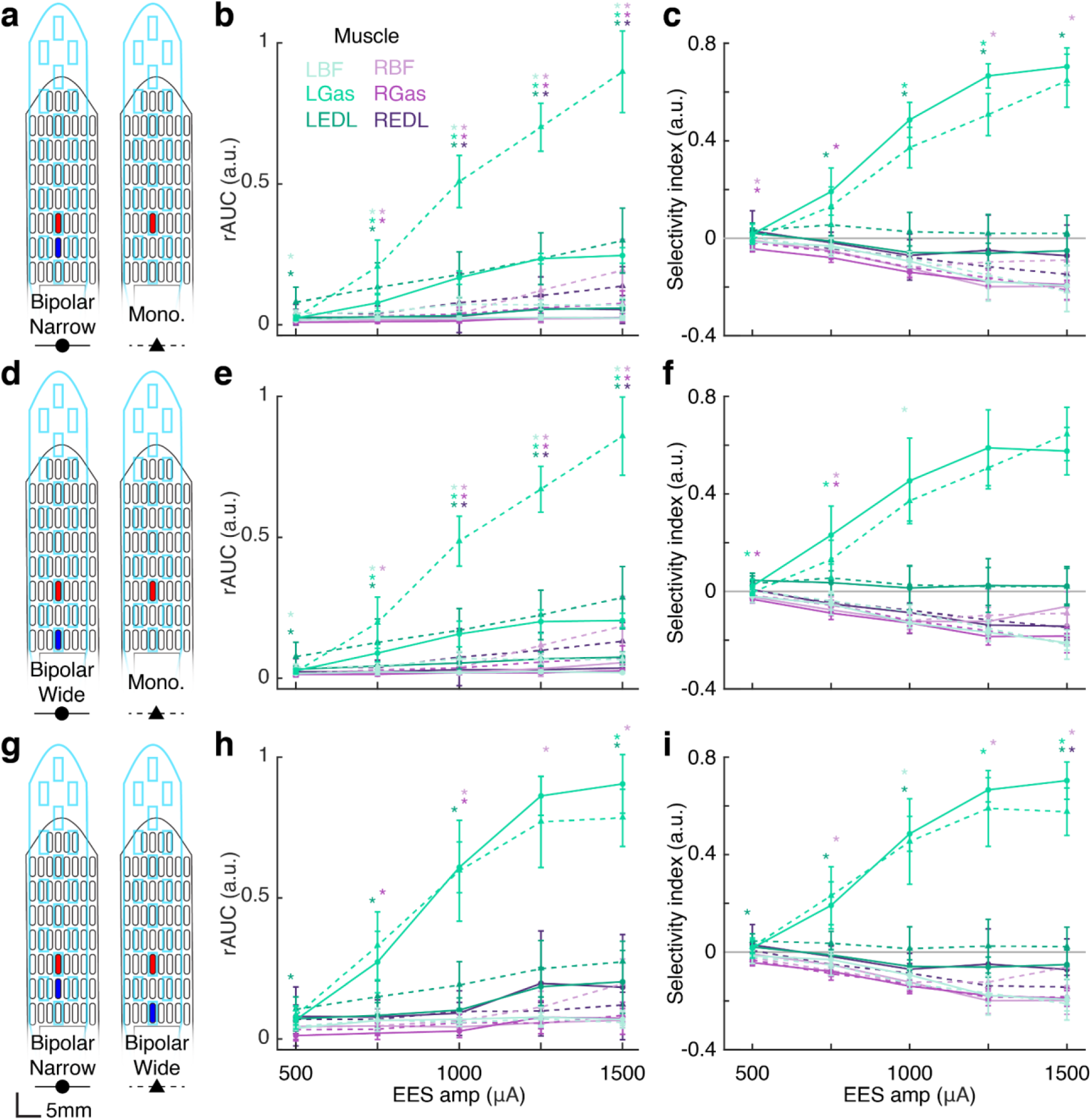
Muscle power and selectivity comparisons in response to bipolar stimulation. **a, d, g)** The stimulation cathode (red) and anode (blue) for the bipolar configurations, and the cathode for the monopolar configurations. In the monopolar configurations, the stimulation current return contact was a wire implanted in the paraspinal muscles. **b, e, h)** Recruitment curves for each stimulation contact configuration presented in the left-hand column. Points are the mean recruitment, and the bars are the standard deviation. **c, f, i)** Selectivity index as a function of amplitude for each stimulation contact configuration presented in the left-hand column. Points are the mean recruitment, and the bars are the standard deviation. Statistical significance was calculated using a Kruskal-Wallis non-parametric test, with a maximum *p*-value of 0.05.

**Supplemental Figure 4.**
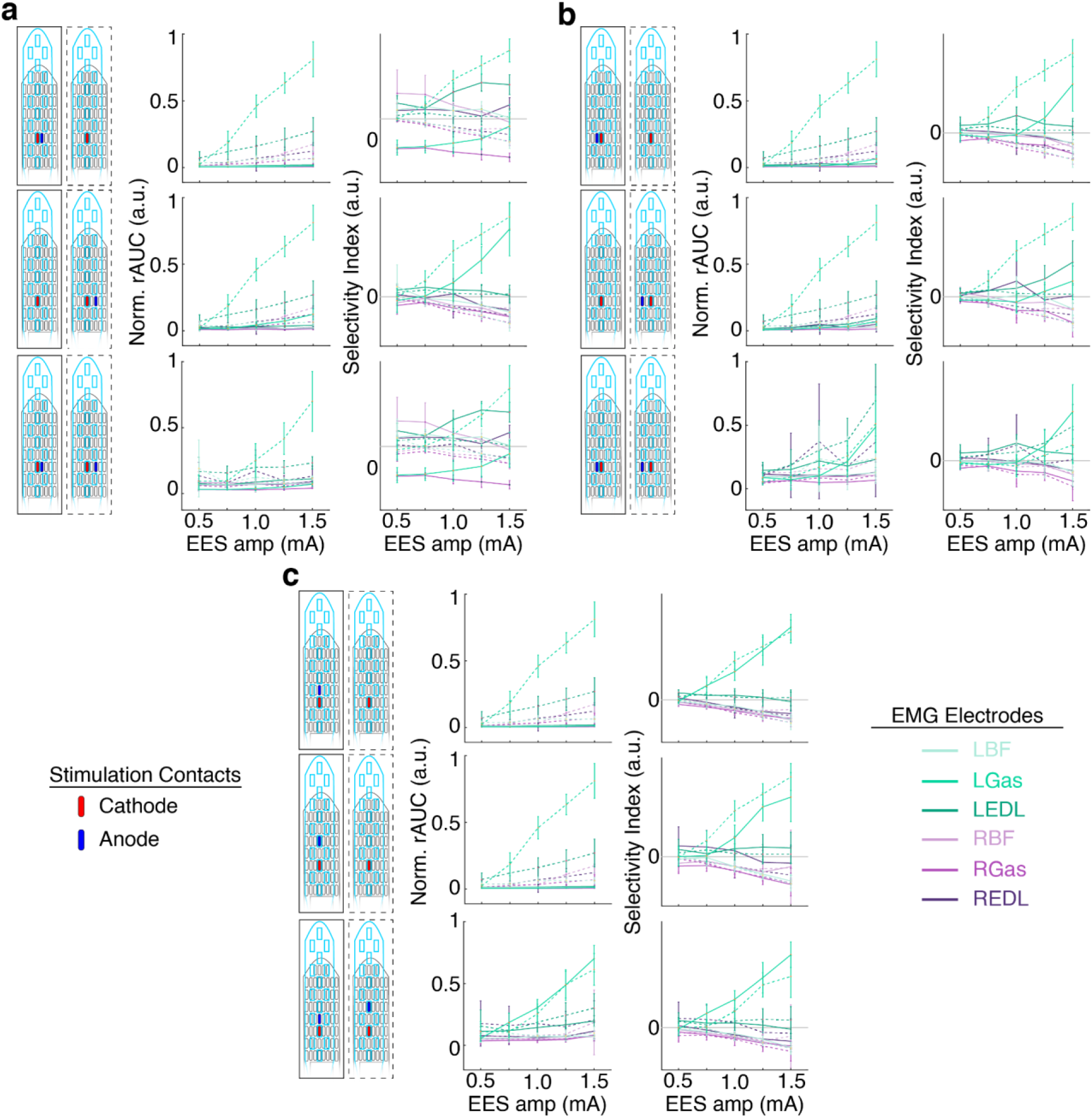
Additional current-steering stimulation-evoked motor response examples. **a-c)** Left column: the stimulation configuration. Red contacts are cathode-leading, blue contacts are anode-leading. An outline of the Medtronic 5-6-5 is shown in blue. The responses to the stimulation shown in the solid border are shown with a solid line in the following columns. In monopolar configurations, the stimulation return was a ground wire implanted in the paraspinal muscles. Middle column: the normalized rectified area under the curve of the electromyography (EMG) responses. Right column: the selectivity indexes of the evoked responses.

**Supplemental Figure 5.**
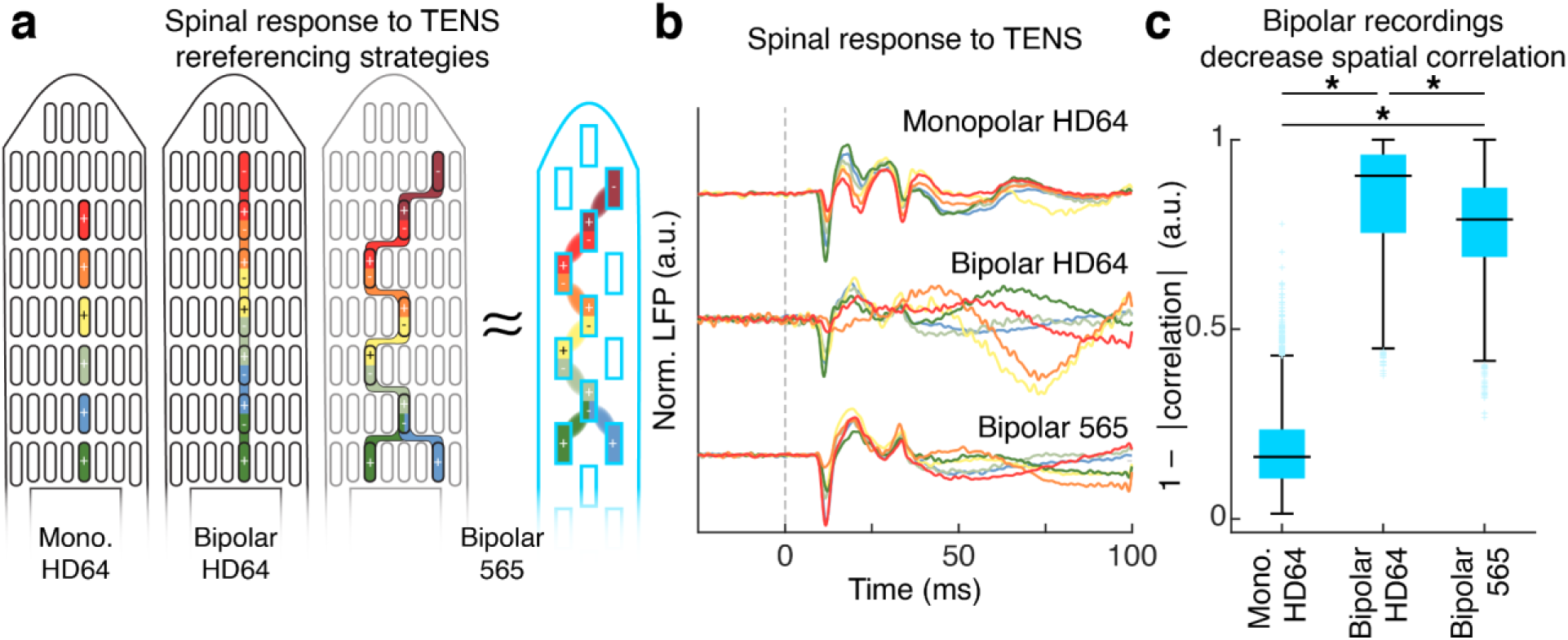
Examining the effect of referencing bipole separation on spatial correlation of peripheral-evoked spinal potentials. **a)** A map of the recording contacts for the three recording configurations. In monopolar configurations, the recording reference was a wire implanted in the epidural space. **b)** The mean sensorimotor evoked potentials (SEPs) recorded from 50 TENS stimulation events (TENS amplitude = 25V) for each of the three recording configurations. **c)** The distribution of spatial uniqueness scores (1 - |correlation|) achieved for each of the three recording configurations. The black horizontal bar is the median, and the boxes extend from the 25th to the 75th percentile. Statistical significance was calculated using a Kruskal-Wallis non-parametric test, with a maximum *p*-value of 0.05.

**Supplemental Figure 6.**
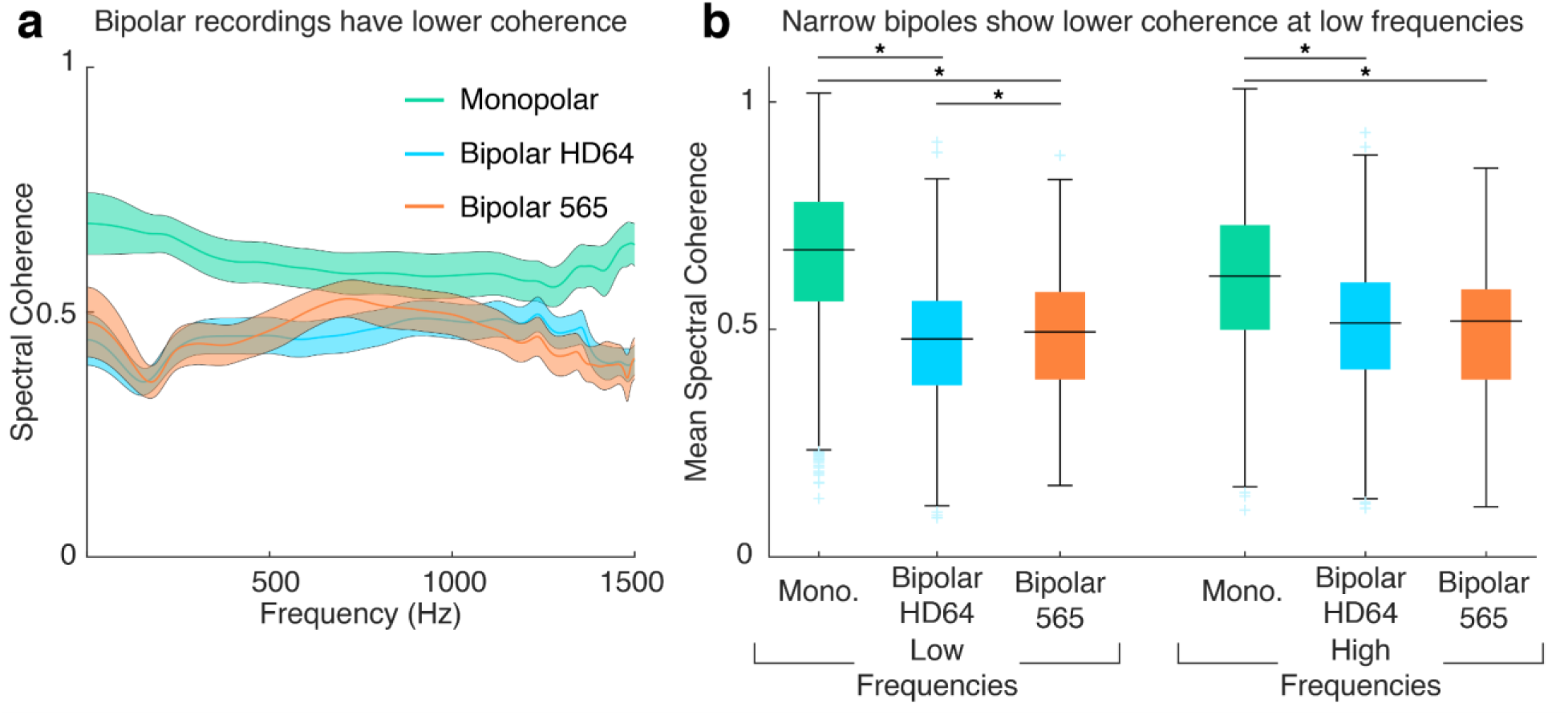
Spectral comparison of spinal local field potentials. **a)** The mean and variance of the pairwise magnitude squared coherence for each channel. High coherence values indicate that all channels record that frequency with equal power. **b)** A boxplot showing mean magnitude squared coherence values. Results are shown for low frequencies (f < 800Hz) and high frequencies (f ≥ 800Hz). Statistical significance was calculated using a Kruskal-Wallis non-parametric test, with a maximum *p*-value of 0.05.

**Supplemental Figure 7.**
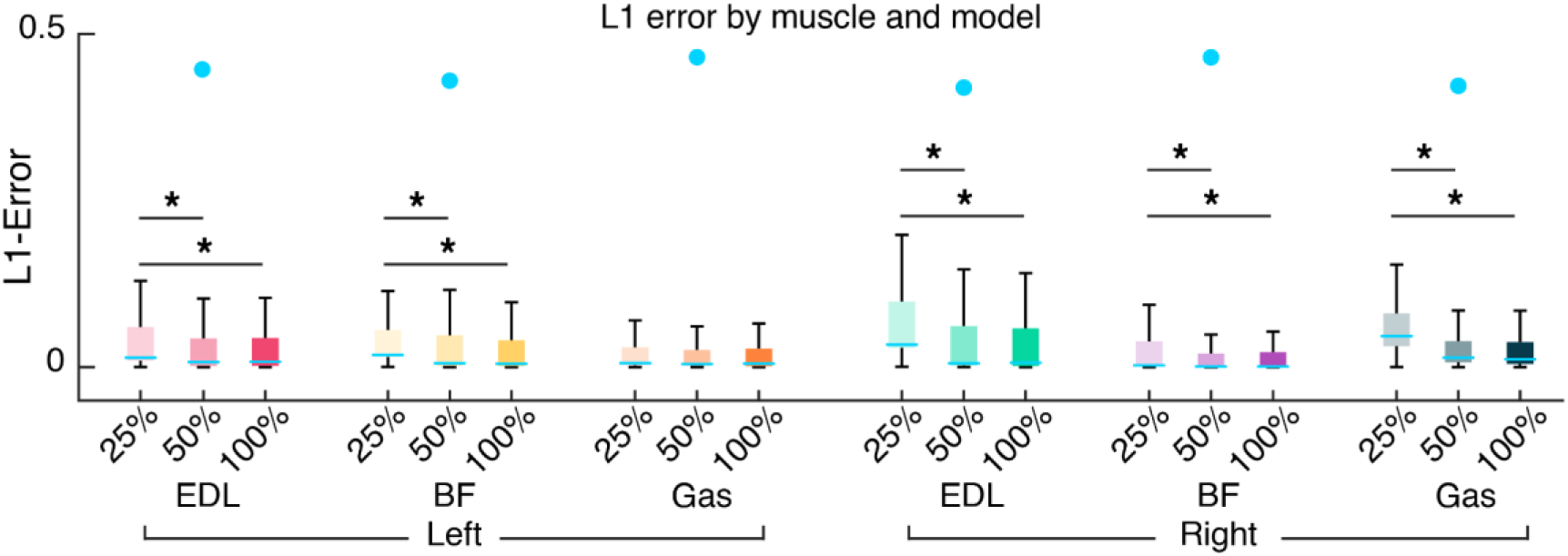
L1 loss by muscle for the 25%, 50% and 100% EES parameter inference neural network models.

**Supplemental Figure 8.**
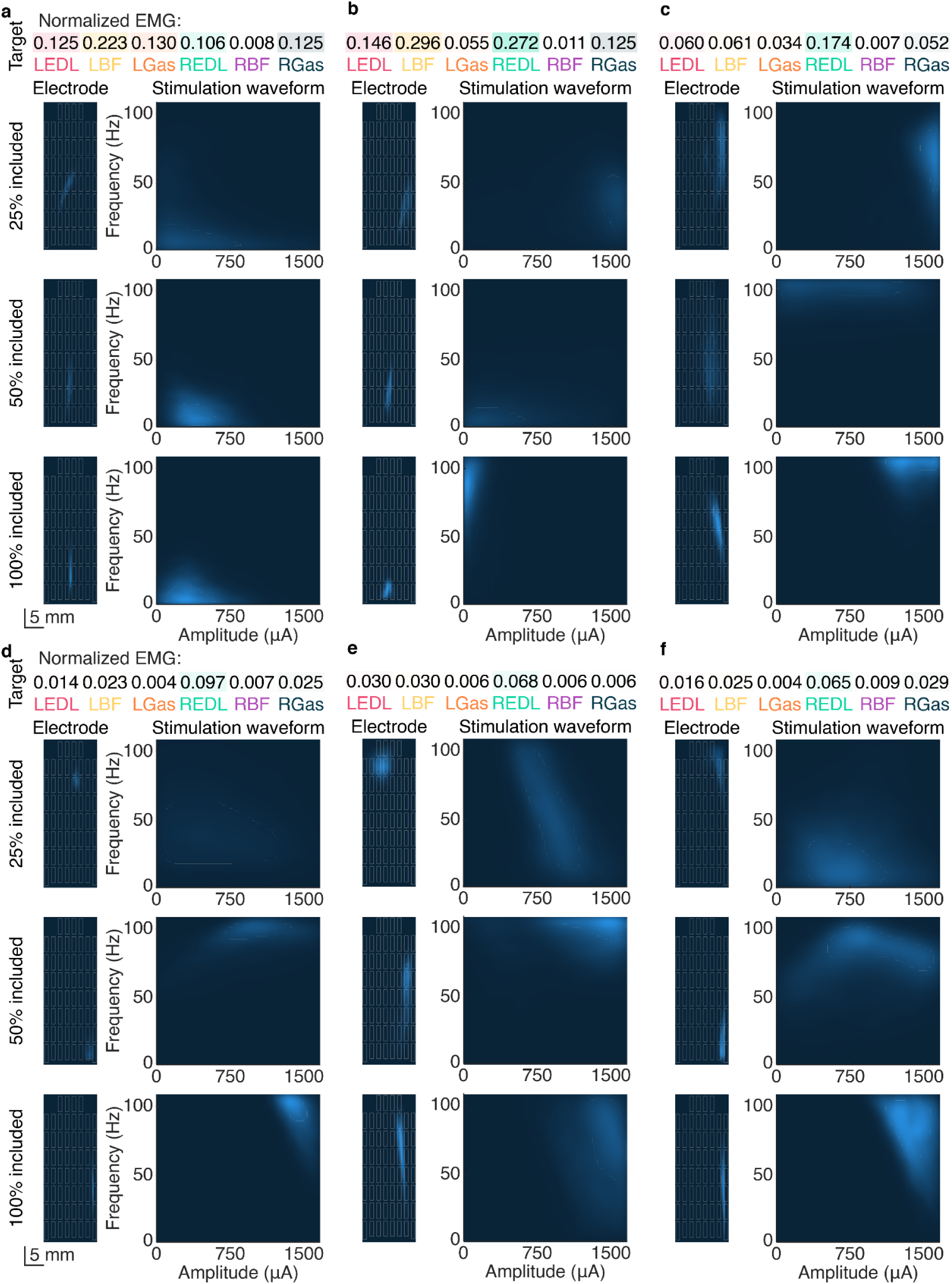
Additional EMG parameter inference posteriors predicted by the inverse model. **a-f)** Posterior likelihood densities for electrode- and amplitude-frequency-space predicted by the 25%, 50% and 100% electrode coverage models. High likelihood regions are shown in lighter colors.

